# Transkingdom mechanism of MAMP generation by chitotriosidase (CHIT1) feeds oligomeric chitin from fungal pathogens and allergens into TLR2-mediated innate immune sensing

**DOI:** 10.1101/2022.02.17.479713

**Authors:** Tzu-Hsuan Chang, Yamel Cardona Gloria, Margareta J. Hellmann, Carsten Leo Greve, Didier Le Roy, Thierry Roger, Francesca Bork, Stefanie Bugl, Johanna Jakob, Lydia Kasper, Bernhard Hube, Stefan Pusch, Neil Gow, Morten Sørlie, Anne Tøndervik, Bruno M. Moerschbacher, Alexander N.R. Weber

**Author notes:** **Contact information (Corresponding Author and Lead Contact)**: Alexander N. R. Weber, Interfaculty Institute for Cell Biology, Department of Immunology, University of Tübingen, Auf der Morgenstelle 15, 72076 Tübingen, Germany.

## Abstract

Chitin is a highly abundant polysaccharide in nature and linked to immune recognition of fungal infections and asthma in humans. Ubiquitous in fungi and insects, chitin is absent in mammals and plants and, thus, represents a microbe-associated molecular pattern (MAMP). However, the highly polymeric chitin is insoluble, which potentially hampers recognition by host immune sensors. In plants, secreted chitinases degrade polymeric chitin into diffusible oligomers, which are ‘fed to’ innate immune receptors and co-receptors. In human and murine immune cells, a similar enzymatic activity was shown for human chitotriosidase (CHIT1) and oligomeric chitin is sensed via an innate immune receptor, Toll-like receptor (TLR) 2. However, a complete system of generating MAMPs from chitin and feeding them into a specific receptor/co-receptor-aided sensing mechanism has remained unknown in mammals. Here, we show that the secreted chitinolytic host enzyme, CHIT1, converts inert polymeric chitin into diffusible oligomers that can be sensed by TLR1-TLR2 co-receptor/receptor heterodimers, a process promoted by the lipopolysaccharide binding protein (LBP) and CD14. Furthermore, we observed that *Chit1* is induced via the β-glucan receptor Dectin-1 upon direct contact of immortalized human macrophages to the fungal pathogen *Candida albicans*, whereas the defined fungal secreted aspartyl proteases, Sap2 and Sap6, from *C. albicans* were able to degrade CHIT1 in vitro. Our study shows the existence of an inducible system of MAMP generation in the human host that enables contact-independent immune activation by diffusible MAMP ligands with striking similarity to the plant kingdom. Moreover, this study highlights CHIT1 as a potential therapeutic target for TLR2-mediated inflammatory processes that are fueled by oligomeric chitin.

## Introduction

Chitin, a hydrophobic polymer of β-1,4-linked N-acetylglucosamine (GlcNAc), is abundant in nature and can be found e.g. in the cell wall of fungi, the exoskeletons of arthropods such as crustaceans and insects, and nematodes (reviewed in [1–3]). However, chitin does not exist in mammals and plants [2, 4, 5]. In these organisms, chitin is a model microbe-associated molecular pattern (MAMP), as evidenced by the existence of chitin-mediated activation of immune responses through pattern recognition receptors (PRRs) [5, 6]. Chitin is sensed through the receptor CERK1 and its co-receptor CEBiP in plants [7] [8] and in humans by Toll-like receptor (TLR) 2 in immune cells [9] and by FIBCD1 or LYSMD3 on epithelial cells [10, 11]. However, during initial exposure, the mammalian host encounters chitin in a highly polymeric and insoluble form of exoskeleton particles of house dust mites or the cell wall of pathogenic fungi, have a form that has been considered immunologically inert [4]. Meanwhile, chitin oligomers of 6 and more GlcNAc subunits have emerged as potent immune-stimulants sensed through PRRs in plants [7] and mammals [9]. This begged two questions. First, how oligomeric chitin subunits are generated in the mammalian host; and second, how the still rather poorly soluble and hydrophobic oligomers are transferred to, and sensed by PRRs?

The generation of MAMPs from complex, polymeric precursors has been observed, for example, in *Drosophila melanogaster* and in plants. In *D. melanogaster*, Gram-negative bacteria-derived binding protein 1 (GNBP1) was reported to hydrolyze Gram-positive peptidoglycan to generate muropeptides [12]. Plant secreted chitinases hydrolyze fungal cell walls to generate chitin oligomers sensed by membrane-expressed chitin receptor complexes (reviewed by [5]). Many plant chitinases belong to the family of glycosyl hydrolases 18 (GH18), a protein family that exists in plants, fungi, bacteria, actinomycetes, insects and humans [13–15]. In addition to acidic mammalian chitinase, humans produce chitotriosidase (CHIT1, also abbreviated as HCHT) [16]. CHIT1 is expressed by neutrophils, macrophages or epithelial cells [17–19] and the dominant chitinase in the human lung [20]. CHIT1 overexpression in inflammatory conditions was first noted for Gaucher disease (MIM #230800), a lysosomal storage disease, where it serves as a therapy biomarker [21]. CHIT1 is also elevated in patients with Type 2 diabetes [22], amyotrophic lateral sclerosis (ALS) [23] [24] [20, 25] and childhood asthma [26], and it is generally accepted that CHIT1 reflects macrophage or microglia activation in these conditions. CHIT1 exists in two major forms: a 50 kDa form mainly found in the blood, and a 39 kDa form expressed predominantly in tissues [27]. Both forms contain a catalytic glycoside hydrolase (GH) 18 domain and display chitinolytic activity. Whereas the full-length, 50 kDa form of CHIT1 contains an additional, C-terminal carbohydrate binding module (CBM) domain, this domain is absent in the 39 kDa form, as it originates from proteolytic cleavage of the C-terminus of the 50 kDa CHIT1 in lysosomes [28]. Because of its ability to break down chitin, chitinase catalytic activity has been thought to contribute to host defense against fungal infections. Indeed, CHIT1 inhibited the growth of *Candida albicans* hyphae, and ectopic expression of CHIT1 in hamster ovary cells restricted the growth of *C. albicans*, *Aspergillus niger* and *Cryptococcus neoformans* [29]. In an in vivo model of systemic candidiasis, Chit1 also contributed to host resistance [19] and genetic chitinase deficiency of *CHIT1* was associated with *C. albicans* colonization in patients with cystic fibrosis [30]. A contribution to macrophage activation via chitin or chitosan degradation had been noted [31]. Conversely, another group observed a significant decrease in kidney fungal burden in *Chit1*-deficient mice compared to mice expressing the functional enzyme. They suggested that CHIT1-generated chitobiose (di-GlcNAc) acted as an immune suppressant [32]. Likewise, in experimental *K. pneumoniae* lung infection, *Chit1*-deficiency limited bacterial dissemination and improved survival, albeit via a different mechanism [33]. Aside from fungal infections, in a murine model of interstitial lung disease, a manifestation of systemic sclerosis, *Chit1*-deficiency prevented fibrotic lung damage while transgenic *Chit1* overexpression promoted it [18]. On the other hand, lack of *Chit1* in an in vivo model of allergic lung inflammation showed reduced induction of protective regulatory T cells and hence a protective role [26]. Depending on the context, CHIT1 thus seems to be able to promote or restrict immune responses. Its integration into the complex system of immuno-stimulation in the host, and its putative role as a MAMP generating enzyme are uncertain.

In addition to MAMP generation from polymeric and hydrophobic precursors, further activities may be required to optimize sensing by mammalian PRRs, like the transfer to and co-engagement of MAMPs at the cell surface. For example, the serum protein lipopolysaccharide binding protein (LBP) and the soluble or membrane glycosyl phosphatidylinositol (GPI)-anchored CD14 both contain hydrophobic binding pockets and transfer lipopolysaccharide (LPS) to TLR4 to initiate signaling [34]. Interestingly, LBP and CD14 also play a role in the transfer of mycobacterial lipopeptides to TLR2 [35, 36]. TLR4 employs the soluble protein MD-2 as part of the actual receptor complex [34], while TLR2 is assisted by the transmembrane co-receptors, TLR1 and TLR6 to sense tri- and di-acetylated mycobacterial lipopeptides, respectively [37, 38]. In plants, the GPI-anchored CEBiP supports the chitin receptor, CERK1 in chitin recognition at the protoplast [39]. However, soluble accessory proteins or co-receptors able to shuttle oligomeric chitin to receptors (e.g. TLR2) have not been described in humans.

In this study, we examined the role of human CHIT1 in chitin sensing. We report that CHIT1 degrades polymeric, relatively inert chitin from *C. albicans* and house dust mites (HDM) to enable TLR2-dependent NF-κB activation and cytokine production. Interestingly, chitin sensing was not strictly dependent on direct contact of TLR2 to chitin but was rather mediated by diffusible oligomers that were generated in the presence of CHIT1. We could further observe that chitin oligomers induced TLR1/TLR2 heterodimerization, and CD14 and LBP facilitated TLR2-NF-κB-dependent chitin sensing. Furthermore, we found that *Chit1* was induced via the β-glucan receptor Dectin-1 in murine macrophages exposed to pathogenic fungal cells, while secreted fungal proteases from *C. albicans* were able to degrade CHIT1 as a potential fungal mechanism of immune escape. Overall, our study unravels a novel and highly conserved system of chitin-based MAMP generation, shuttling and recognition in mammals.

## Results

### CHIT1 generates diffusible TLR2-activating oligomeric chitin ligands

Based on the reported chitinase-based MAMP-generating system in plants [5], we first sought to investigate whether circulating enzymes that generate MAMPs for specific PRRs might play a role in chitin sensing in humans, e.g. for TLR2 activator [9]. Since acidic mammalian chitinase displays exochitinase activity [40], it would be expected to release only di-GlcNAc subunits which are too small to stimulate TLR2 activator [9]. However, CHIT1 is an endochitinase, cleaving inside chitin chains as opposed to [17, 41]. Thus, CHIT1 could enable the generation of longer oligomeric chitin fragments [31] that might directly stimulate TLR2. To test this hypothesis, purified commercially available macroscopic chitin flakes from shrimp were digested with purified recombinant 39 kDa (lacking the CBD) or 50 kDa (containing the CBD) recombinant CHIT1 (see [42] and Methods) and transferred the product-containing reaction to TLR2-expressing NF-κB reporter HEK293T cells (HEK293T-TLR2). Interestingly, while macroscopic shrimp chitin was unable to activate TLR2, incubation with 39 kDa CHIT1 rendered it TLR2-active (Fig. 1A). A similar TLR2 stimulation was observed when the experiment was repeated using a transwell setup separating the chitin flake-containing degradation reaction from the cells (Fig. 1B). By doing so, we excluded the possibility that TLR2-stimulating activity might be mediated by direct contact of cells, and hence TLR2, with macroscopic chitin (size approximately 0.5 × 1-2 mm, Fig. S1A), which was retained by the transwell insert (Fig. S1B). Conversely, lower molecular weight chitin oligomers (estimated molecular weight 2000-3000, size ∼5 nm, [9]) were able to pass the filter pores and reach the cells. Although recombinant CHIT1 was clearly not able to activate TLR2 by itself, we further ruled out a possible role of confounding contaminants in recombinant CHIT1 protein preparation via ectopically expressing CHIT1 in HEK293T. In transfected cells, both CHIT1 isoforms were secreted (see Fig. S1C for immunoblots of cell SNs) and incubation of CHIT1-containing cell culture media also rendered the otherwise inactive macroscopic chitin TLR2-active (Fig. 1C). To demonstrate further that activation by CHIT1 was truly due to enzymatic degradation of macroscopic chitin, we mutated the catalytic site of CHIT1 (D138A E140L, *cf.* [43], Fig. S1D). Medium from cells transfected with a mutant CHIT1 (mCHIT1) plasmid did contain secreted mCHIT1 (Fig. S1D) but, as expected, lacked chitinase activity (Fig. S1E) and was unable to elicit TLR2 activity (Fig. 1D). We conclude that that CHIT1 was able to generate diffusible TLR2 activating ligands from macroscopic shrimp chitin.

**Figure 1:**
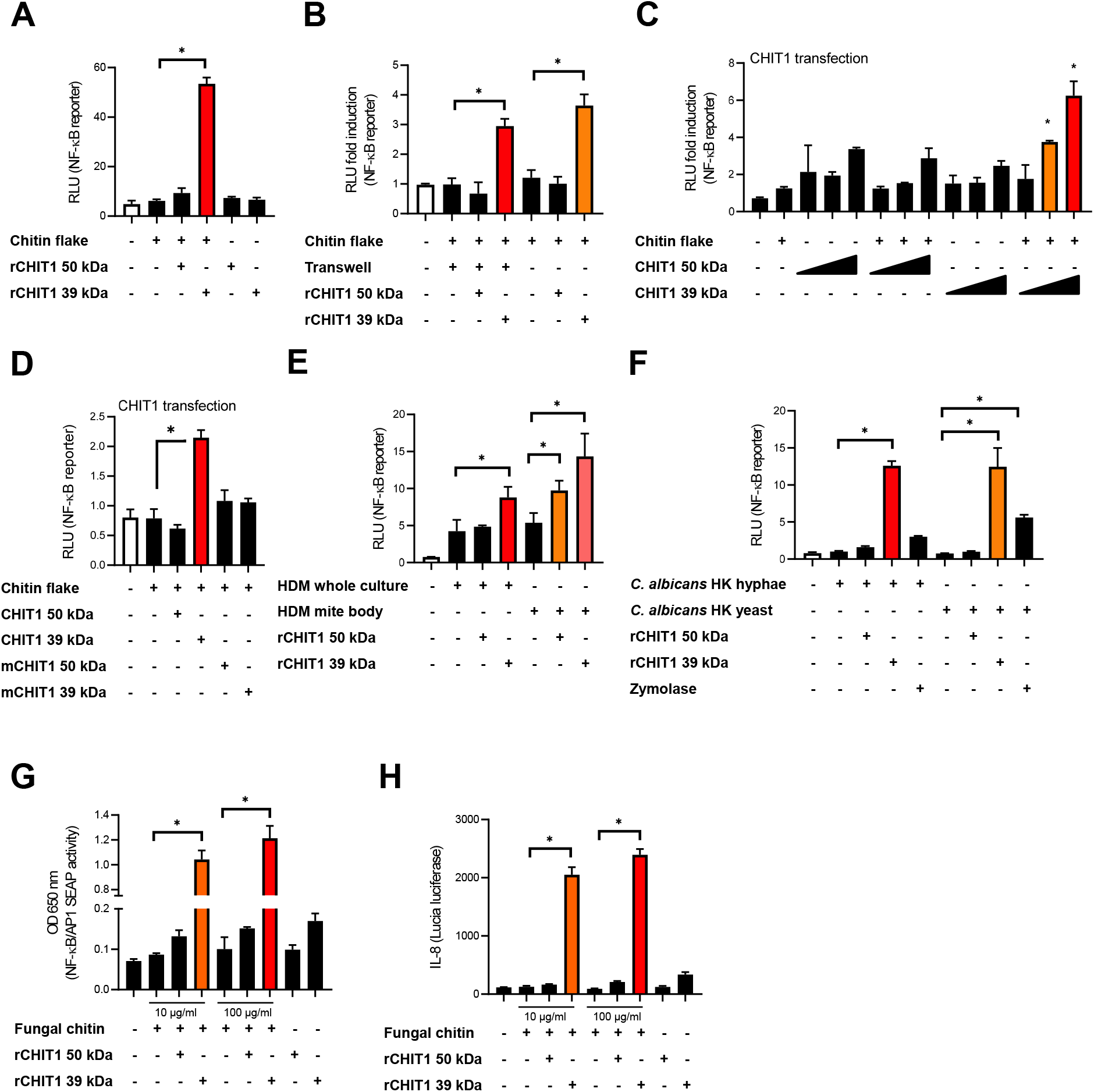
CHIT1 converts poorly immunogenic polymeric chitin into TLR2-active chitin. (A-H) CHIT1 digests of macroscopic chitin induce NF-κB activity in TLR2-overexpressing HEK293T cells. Recombinant CHIT1 isoforms (rCHIT1, in A, B, E-H) or supernatants from HEK293T cells expressing either wild-type CHIT1 (in C and D) and/or catalytically-inactive CHIT1 (mCHIT1) plasmids (in D) were incubated with chitin flakes (A-D), house dust mite (HDM) whole culture or mite bodies (E), *Candida albicans* hyphae and yeast forms (F) or highly purified *C. albicans* fungal chitin (G and H). After 18 h, the CHIT1-incubated culture supernatants were transferred to TLR2- and NF-κB-transfected HEK293T cells (A-F) or HEK-Dual™ hTLR2 reporter cells (G, H) for 18 h and lysates used for triplicate dual luciferase assays (A-F) (n = 3) or supernatants for triplicate NF-κB (G) or *IL8* (H) promotor reporter analysis. In (B) the transwell setting was applied to avoid direct contact of undigested (−) and rCHIT1-digested (+) chitin flakes with the cells at the bottom of the plate. In A (n=3), C (n=3), D n=4), E (n=2), F (n=3), G and H (n=4 each) one representative of ‘n’ biological replicates are shown (mean+SD for technical replicates). B (n=2) represents combined data (mean+SD) from ‘n’ biological replicates. * p<0.05 according to one-way ANOVA with Dunnett’s (A), Tukey’s (B, D and E) or Sidak’s (C, F, G and H) correction for multiple testing.

We then went on to investigate whether this effect applied to polymeric chitin from other sources, such as the disease-relevant HDM, *Dermatophagoides pteronyssinus* and the pathogenic fungus, *C. albicans*. Using the same assay, we observed a significant increase of TLR2 activity upon exposure of HEK293T-TLR2 to *D. pteronyssinus* whole cultures and mite bodies pre-incubated with recombinant 39 kDa CHIT1 in both the normal and transwell setups (Fig. 1E). The same effect was observed for chitin derived from heat-killed *C. albicans* yeast and hyphae (Fig. 1F). In this experiment, zymolase, an enzyme cleaving β-glucan chains, only had a minor effect that may be attributable to degrading the scaffolding to which chitin is attached in the cell wall. The experiment was also performed specifically with macromolecular, highly purified *C. albicans* chitin [44], which triggered a drastic increase in TLR2-mediated activity upon 39 kDa CHIT1 digestion (Fig. 1G and H). Collectively, these data show that CHIT1 is able to generate potent and diffusible TLR2-activating MAMPs from polymeric, poorly activating macroscopic chitin, reminiscent of observations done in *Drosophila* [12] and plants [5].

### CHIT1 binds to the fungal cell wall

To verify the effect of CHIT1 on fungal cell wall chitin, we performed confocal fluorescence microscopy. *C. albicans* yeast cells were treated with caspofungin, an inhibitor of β-(1,3)-D - glucan synthesis, that increases chitin exposure on the fungal cell wall [45]. Caspofungin-treated cells were heat-inactivated and pre-incubated with 39 kDa or 50kDa CHIT1. Finally, yeast cells were either labelled with the fluorescein-conjugated, chitin-binding lectin wheat-germ agglutinin (WGA) or fluorescein-conjugated concanavalin (Con) A, which binds to α-mannans. Confocal imaging revealed that WGA bound prominently to *C. albicans* yeast cells upon CHIT1 digestion, whereas binding of ConA remained unchanged (Fig. 2A, quantified in 2B). Additionally, 50 kDa CHIT1 stably bound to and could be visualized at the fungal cell wall (Fig. 2C). No detectable binding of the 39 kDa form was observed, probably due to the lack of a CBD and hence only transient interaction with chitin [42]. Its lower retention time at the cell wall would explain the generation of longer soluble oligomers, which are more favorable TLR2 ligands [9], by the 39 kDa form due to lower processivity. In summary, CHIT1 can directly engage fungal cell wallw, consistent with the release of oligomeric chitin fragments.

**Figure 2.**
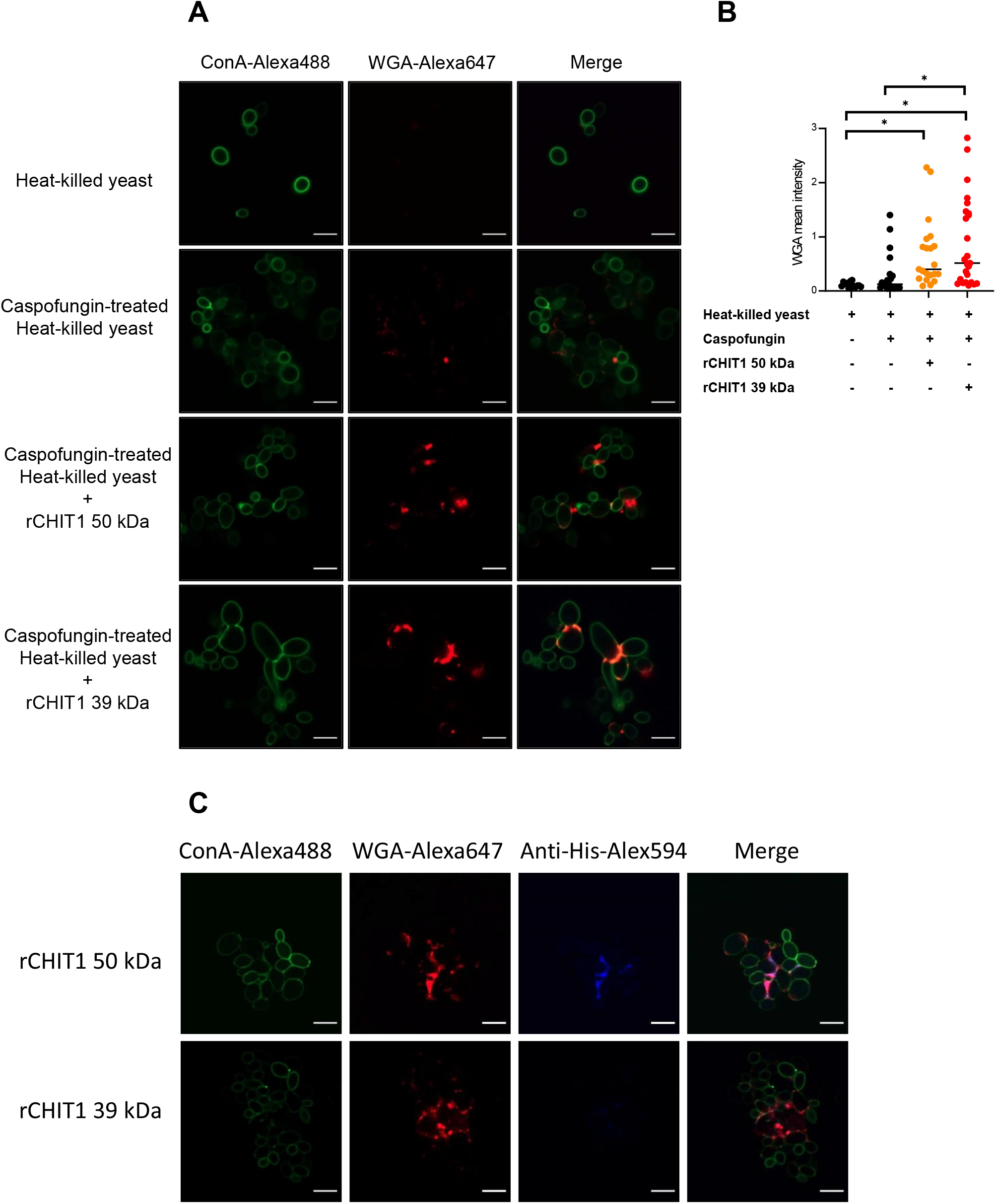
CHIT1 digestion increases chitin exposure on *C. albicans* yeast. (A, C) Heat-killed *C. albicans* yeast cells were left untreated or treated with caspofungin (0.032 μg/ml) and CHIT1 as indicated. Cells were then stained with Alexa448-ConA (green) and Alexa647-WGA (red). Shown are representative single focal images of *C. albicans* yeast cells containing at least 5-20 yeast cells, with multiple images taken per condition across multiple biological replicates. (B) WGA fluorescent intensity was quantified, thresholds applied to the ConA and WGA signals and the intensity measured by using the Otsu method. The mean intensity of WGA signal was normalized to that of ConA. (C) To stain CHIT1 on caspofungin-treated, heat-killed *C. albicans* yeast, Alexa594-conjugated anti-His antibodies were used to detect the His-tag on CHIT1. In A (n=3) and C (n=2) one representative of ‘n’ biological replicates is shown. B (n=2) represents combined data (mean and individual points) from ‘n’ biological replicates (each dot represents one quantified image). * p<0.05 according to one-way ANOVA with Tukey’s correction for multiple testing.

### Chitin binding involves multiple PRRs, whereas TLR2 can sense diffusible oligomeric chitin

The TLR2-HEK293T system lacks many PRRs except TLR2 and TLR2 co-receptors [46]. To extend our insights to a more relevant system, bone marrow-derived macrophages (BMDM) were stimulated with native and CHIT1-digested *D. pteronyssinus* and *C. albicans* hyphae and yeast. BMDM produced IL-6 and TNF in response to undigested *D. pteronyssinus* and *C. albicans*. However, the stimulatory capacity of HDM (Fig. 3A-B), yeast and hyphae (Fig. 3C-D) increased following CHIT1 39 kDa digestion, albeit mainly for IL-6. Analysis of *Tlr2* KO BMDM and use of the transwell setup showed that both TNF and IL-6 were induced by HDM preparations primarily upon direct contact in a strongly TLR2-dependent manner (Fig. 3E and F). Likewise, In contrast *C. albicans* hyphae stimulated TNF and IL-6 production mainly upon direct contact. Yet with chitin, diffusible components induced IL-6 secretion in a TLR2-dependent manner as evidenced using transwell experiments (Fig. 3G and H). This suggests that in BMDM and for *C. albicans,* TLR2 is not only able to elicit cytokine responses to chitin-containing pathogens upon direct binding but appears to be a sentinel for diffusible TLR2 activating ligands generated by CHIT1. Thus, CHIT1 digestion enables PRR-expressing cells to overcome the requirement of direct physical contact with chitin-bearing pathogens by using oligomer- and diffusion-mediated distal chitin sensing.

**Figure 3.**
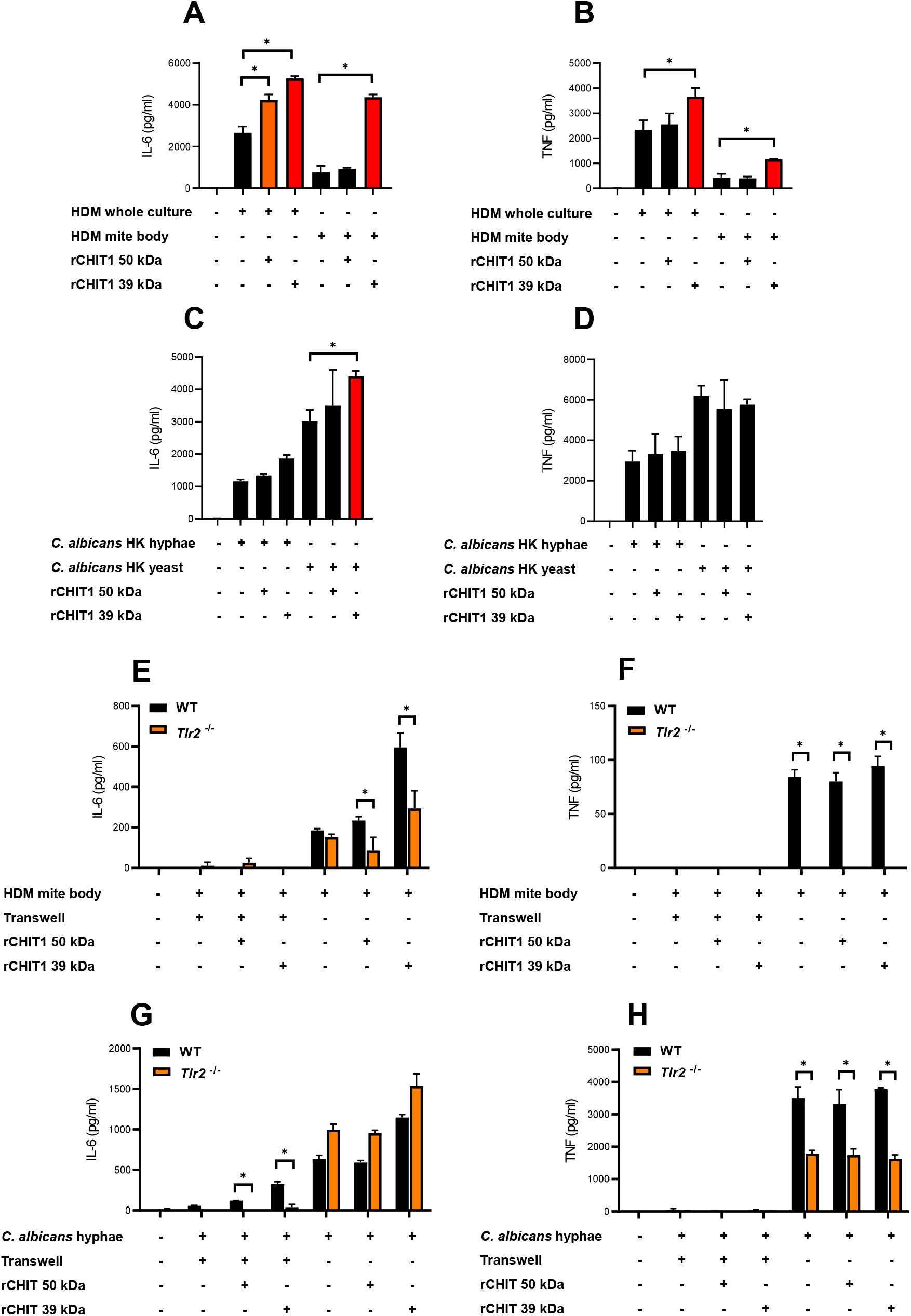
Diffusible chitin oligomers released from chitin-rich organisms elicit the secretion of TLR2-dependent pro-inflammatory cytokines in murine macrophages. (A-H) IL-6 and TNF secretion in WT or *Tlr2* KO murine BMDMs upon stimulation with the indicated organisms left undigested (−) or digested with either isoform of recombinant CHIT1 (+) as assessed via ELISA. (E-H) The culture supernatants of the indicated organisms were digested with CHIT1 and then used to stimulate BMDMs in a transwell setting. IL-6 and TNF secretion was then assessed via ELISA. In A-D (n=2) and E-H (n=3) one representative of ‘n’ biological repeats is shown (mean+SD for technical triplicates). * p<0.05 according to two-way ANOVA with Sidak’s correction for multiple testing.

### Oligomeric chitin signals via LBP, CD14 and TLR1/TLR2 heterodimer

A feature of certain innate immune signaling pathways is the involvement of soluble and membrane-bound transfer proteins such as - LBP and CD14 that mediate LPS sensing by TLR4 and Pam_2_CSK_4_ (Pam2) and Pam3_3_CSK_4_ (Pam3) lipopeptides by TLR2/TLR6 and TLR1/TLR2, respectively [34, 35, 47]). To test whether LBP and CD14 play a role in shuttling oligomeric DP10-15 chitin (C10-15, see [9]), we added recombinant LBP exogenously to and co-expressed CD14 in TLR2-HEK293T cells before measuring NF-κB reporter activity. Evidently, CD14 overexpression strongly increased TLR2-mediated NF-κB activity, and this was further increased by addition of LBP like for Pam3 (Fig. 4A and Fig. S2), indicating that oligomeric chitin sensing employs components of other TLR ligand sensing systems in this recognition pathway. We next tested whether oligomeric chitin TLR2 sensing required co-receptors such as TLR1 and TLR6 [48] by using BMDM from *Tlr1* knockout (-/-, KO), *Tlr2* KO or *Tlr6* KO mice. Interestingly, in response to the oligomeric chitin, C10-15, and to Pam3, *Tlr6* KO BMDM responded like WT BMDMs by producing IL-6 and TNF, whereas both *Tlr1* KO and *Tlr2* KO BMDMs were reduced in their responsiveness to C10-15 and Pam3 (Fig. 4B-C and S3). This genetic analysis implicating both TLR1 and TLR2 in chitin sensing of immune cells was consistent with the earlier use of blocking antibodies directed against TLR1 or TLR6 in HEK293T cells [9]. To validate that oligomeric chitin also triggered receptor dimerization as shown for TLR4 and TLR2, we conducted fluorescence complementation assays of TLR2 with the known TLR2 co-receptors TLR1 or TLR6. In brief, we used expression constructs in which the TLR cytoplasmic domain is C-terminally fused to either the N- or C-terminal portion of the mLumin protein, a derivative of the far-red fluorescent protein variant of mKate [49]. In this system, stable dimer formation is evidenced by the detection of mLumin fluorescence [49]. C10-15 and Pam3 induced the highest mLumin fluorescence in TLR1- and TLR2-expressing cells (Fig. 4D), and Pam2 did so in TLR2- and TLR6-expressing cells (Fig. 4E). Quantification of these data showed that TLR1/TLR2 co-expression induced significant mLumin fluorescence only upon stimulation with C10-15 oligomeric chitin or Pam3 (Fig. 4F), whereas TLR2/TLR6 co-expression did so only upon stimulation with Pam2 (Fig. 4G). Similar results were obtained for NF-κB induction in cells expressing TLR2 and co-receptors (Fig. 4H). Additionally, we found that certain cell types, for example N/TERT-1 keratinocyte-like cells [50], naturally responded to Pam2, whereas they failed to respond to both C10-15 and Pam3 (Fig. 4I), indicating the latter agonists behave similarly and both prefer TLR1/TLR2 heterodimers. This is consistent with *in silico* analyses, which showed that a chitin decamer can be fitted into the structure of a TLR1-TLR2 heterodimer (PDB: 2Z7X, [37], Fig. 4J), whereas the absence of a hydrophobic pocket in TLR6 prevented this in a TLR2-TLR6 heterodimer structure (PDB: 3A79, [38]). Collectively, our results show that oligomeric chitin sensing does not only rely on the generation of oligomeric MAMPs by CHIT1, but also on soluble and membrane-bound co-receptors such as CD14 and TLR1, for optimal TLR2 signal induction.

**Figure 4:**
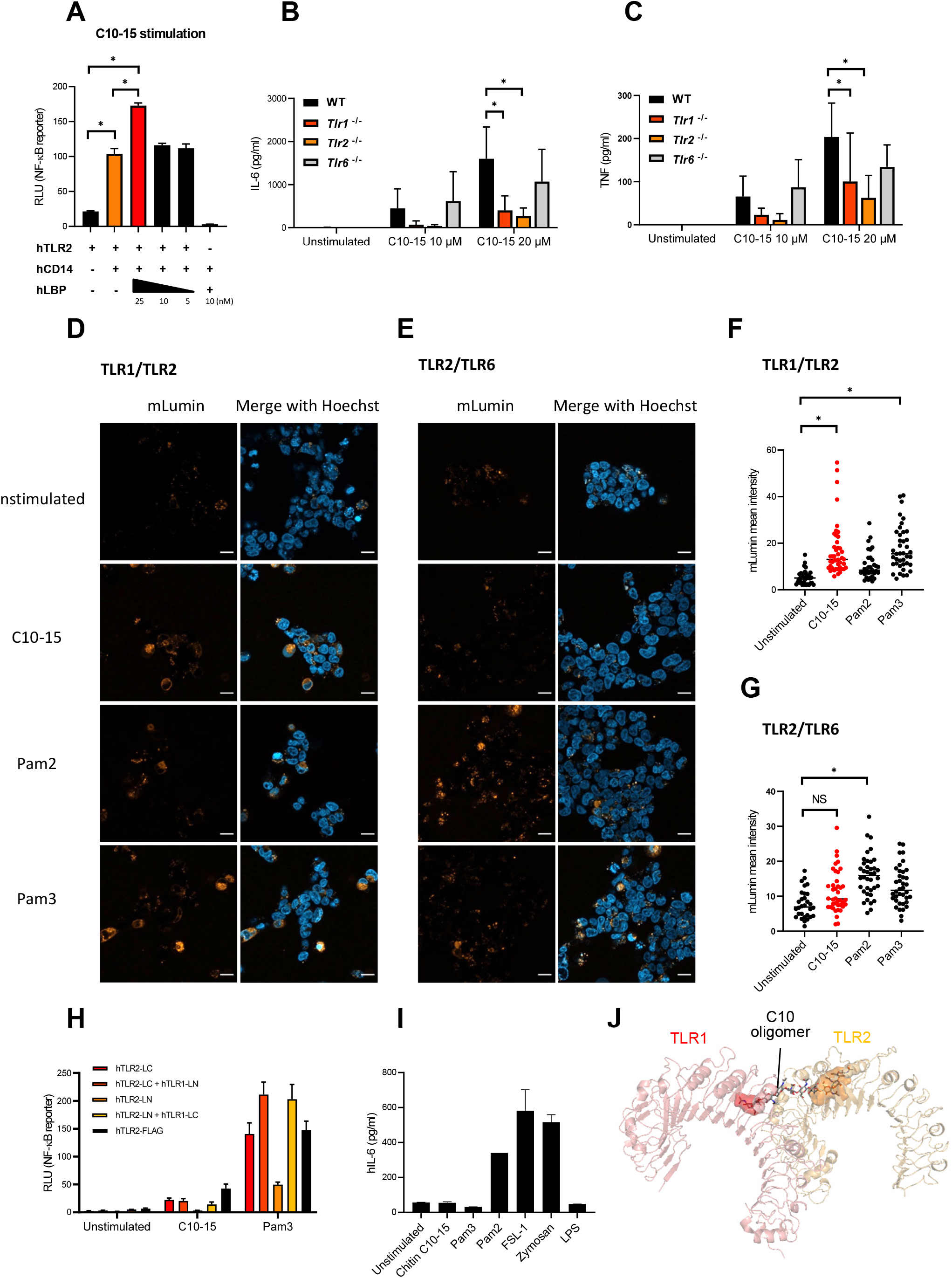
LBP, CD14 and TLR1, but not TLR6, cooperate with TLR2 to mediate oligomeric chitin recognition. (A) Measurement of NF-κB activity in TLR2- or TLR2/CD14-overexpressing HEK293T cells. Dose titrated recombinant LBP protein was added together with chitin C10-15 oligomers. Cell lysates were used for dual luciferase assays done in technical triplicates. (C-H). (B, C) IL-6 and TNF secretion in murine WT, *Tlr1* KO, *Tlr2* KO and *Tlr6* KO BMDMs upon stimulation with chitin C10-15 was measured via triplicate ELISA. (D, E) HEK293T cells were co-transfected with split-mLumin-TLR1 and -TLR2 (D) or split-mLumin-TLR2 and -TLR6 (E). After stimulation with Pam2, Pam3 or C10-15, cells were stained with Hoechst and inspected by confocal fluorescence microscopy. Single representative focal images selected from at least 20 – 25 of cells were taken per condition. Scale bars represent 20 µm. (F, G) The mLumin fluorescent intensity was quantified by ImageJ. Threshold was applied to the fluorescent signal from each single cell and the intensity measured by the Otsu method. The mean intensity of mLumin was normalized to that of Hoechst. (H) Measurement of NF-κB activity in split-mLumin single- or co-transfected HEK293T cells after stimulation with C10-15 or Pam3. (I) IL-6 secretion in human N/TERT-1 cells upon stimulation with the indicated ligands was measured by triplicate ELISA. (J) Fitting of chitin decamer (sticks) into the TLR1 (red)-TLR2 (orange) heterodimer (cartoon, hydrophobic pocket shown as surface) based on crystal structure (PDB: 2Z7X). In A, D, E and I (n=3 each) and H (n=1 preliminary experiment) one representative of ‘n’ biological replicates is shown (mean+SD for technical replicates). B, C (n=4 each), F and G (n=3 each) represent combined data (mean+SD) from ‘n’ biological replicates (in F and G each dot represents one quantified image). * p<0.05 according to one-(A, F, G) or two-way (B, C) ANOVA with Sidak’s (A-C), Tukey’s (F, G) correction for multiple testing.

### Induction of *Chit1* expression by fungal components via Dectin-1

Our data thus far suggest that CHIT1 can generate diffusible, TLR2-activating oligomers from chitin-containing organisms. In murine BMDMs, RT-qPCR analyses revealed a relatively low expression of *Chit1* compared to the housekeeping gene *Tbp* (encoding the TATA box binding protein). Exposure to already oligomeric chitin (C10-15) or to Pam3 did not increase *Chit1* transcription (Fig. 5A), neither did HDM bodies (Fig. 5B). However, *C. albicans* hyphae induced significantly more than two-fold *Chit1* mRNA levels after 2 h stimulation (Fig. 5C). The effect was transient and returned to baseline within 6 h. To test whether this induction was TLR2-dependent, WT and *Tlr2* KO BMDM were compared. Whereas Pam3, C10-15 and *C. albicans* hyphae elicited a strong, *Tlr2*-dependent *Il6* upregulation, *C. albicans* hyphae upregulated *Chit1* and *Il6* in a *Tlr2*-independent manner (Fig. 5D, E). Thus, in murine macrophages rapid induction of *Chit1* was independent of TLR2, and also of chitin. These results are consistent with the fact that, in fungal cell walls, chitin is buried under layers cell wall components including β-glucan and mannoproteins [1, 51]. At first encounter with a fungal pathogen, there would thus be little accessible chitin, and an induction of chitinase activity via another MAMP such as β-glucan would, hence, be more plausible. We, therefore, speculated that *Chit1* induction by *C. albicans* – which is consistent with earlier analyses using macromolecular chitin in primary macrophages [9] – could be mediated by β-glucan sensing through Dectin-1 (encoded by *Clec7A*, [52]). We compared the expression of *Chit1* and *Il6* in WT and *Clec7a* KO immortalized macrophages [9, 53] by RT-qPCR analysis upon stimulation with *C. albicans*, Pam3, C10-15 and zymosan, a yeast cell wall extract sensed through TLR2 and Dectin-1 [9]. Interestingly, soluble TLR2 agonists (Pam3, C10-15) did not show differences between cell types for *Chit1* and *Il6* transcription (Fig. 5F, G). However, *C. albicans* and zymosan induced drastically less *Chit1* and *Il6* by *Clec7a* KO macrophages (Fig. 5F, G). These data indicate that Dectin-1 might be responsible for inducing the expression of *Chit1* in response to *C. albicans*. This would be consistent with β-glucan being a major and more readily exposed fungal MAMP [52].

**Figure 5.**
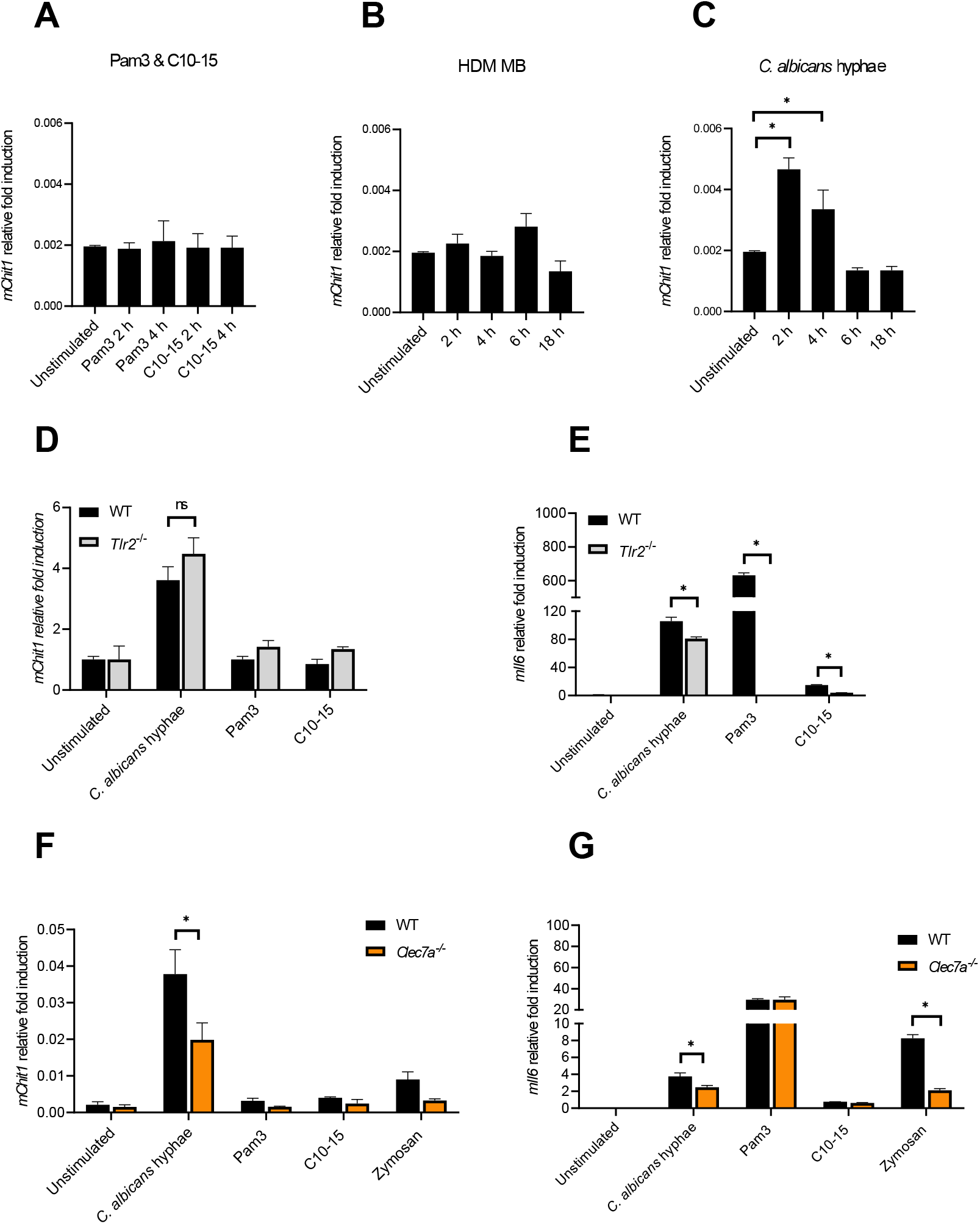
*C. albicans* hyphae elicit *Chit1* mRNA induction via Dectin-1. (A, B and C) Fold induction of murine *Chit1* and *Il6* mRNA relative to *Tbp* mRNA in primary BMDMs upon stimulation with HDM, *C. albicans* hyphae, Pam3 and C10-15 at the indicated time points. (D-G) Relative fold induction of murine *Chit1* and *Il6* mRNA relative to *Tbp* mRNA in (D and E) WT and *Tlr2* KO primary BMDMs or (F and G) WT and *Clec7a*^−/−^ iMacs upon 4 h stimulation with *C. albicans* hyphae, Pam3, C10-15 and zymosan (n = 3). In A-C (n=4), D-G (n=2) one representative of ‘n’ biological replicates is shown (mean+SD for technical replicates). * p<0.05 according to one-way ANOVA with Dunnett’s correction (A-C) or two-way ANOVA with Sidak’s correction (D-G) for multiple testing, respectively.

### CHIT1 is degraded by the fungal proteases Sap2 and Sap6

Based on these results, increasing Chit1 activity and the subsequent emergence of diffusible TLR2 ligands during infection over time would mean an increasing contribution of TLR2-dependent immunity to antifungal responses. This would make CHIT1 a potential target for fungal counter-strategies. Indeed, in plants, chitinases have been shown to represent proteolytic targets of fungal proteases [5]. Since *C. albicans* can express multiple secreted aspartyl proteinases (Saps) [54], we tested whether *C. albicans* culture supernatants contained proteolytic activity to which CHIT1 might be sensitive. We incubated (His-tagged) recombinant Chit1 preparations with fungal supernatants from *C. albicans* grown under conditions known to induce Sap protease secretion (in ‘induction media’ [55]). This led to a decrease of anti-His signal, i.e. CHIT1 protein levels (Fig. 6A), when compared to samples without *C. albicans* supernatant (lanes 9 and 10 in Fig. 6A). Degradation was not observed when *C. albicans* supernatant was harvested from fungal cultures that had been grown in standard YPD media, where Sap proteases are only expressed at low levels [55]. This was confirmed by quantification analysis from several experiments (Fig. 6C and D). These results indicate under these given conditions *C. albicans* cells might secrete Sap proteases to target Chit1. To test this hypothesis further, Chit1 was exposed to purified recombinant *C. albicans* Sap2 and Sap6, representing Sap proteases expressed by yeast or hyphae. Both 50 kDa and 39 kDa forms of CHIT1 were dose-dependently degraded by Sap2 and Sap6 (Fig. 6E). The effect was blocked by the aspartic protease inhibitor pepstatin A (Fig. 6F). Thus, Chit1 is sensitive to the fungal virulence factors and proteases Sap2 and Sap6 and might therefore be targeted by the pathogenic fungus, *C. albicans*.

**Figure 6.**
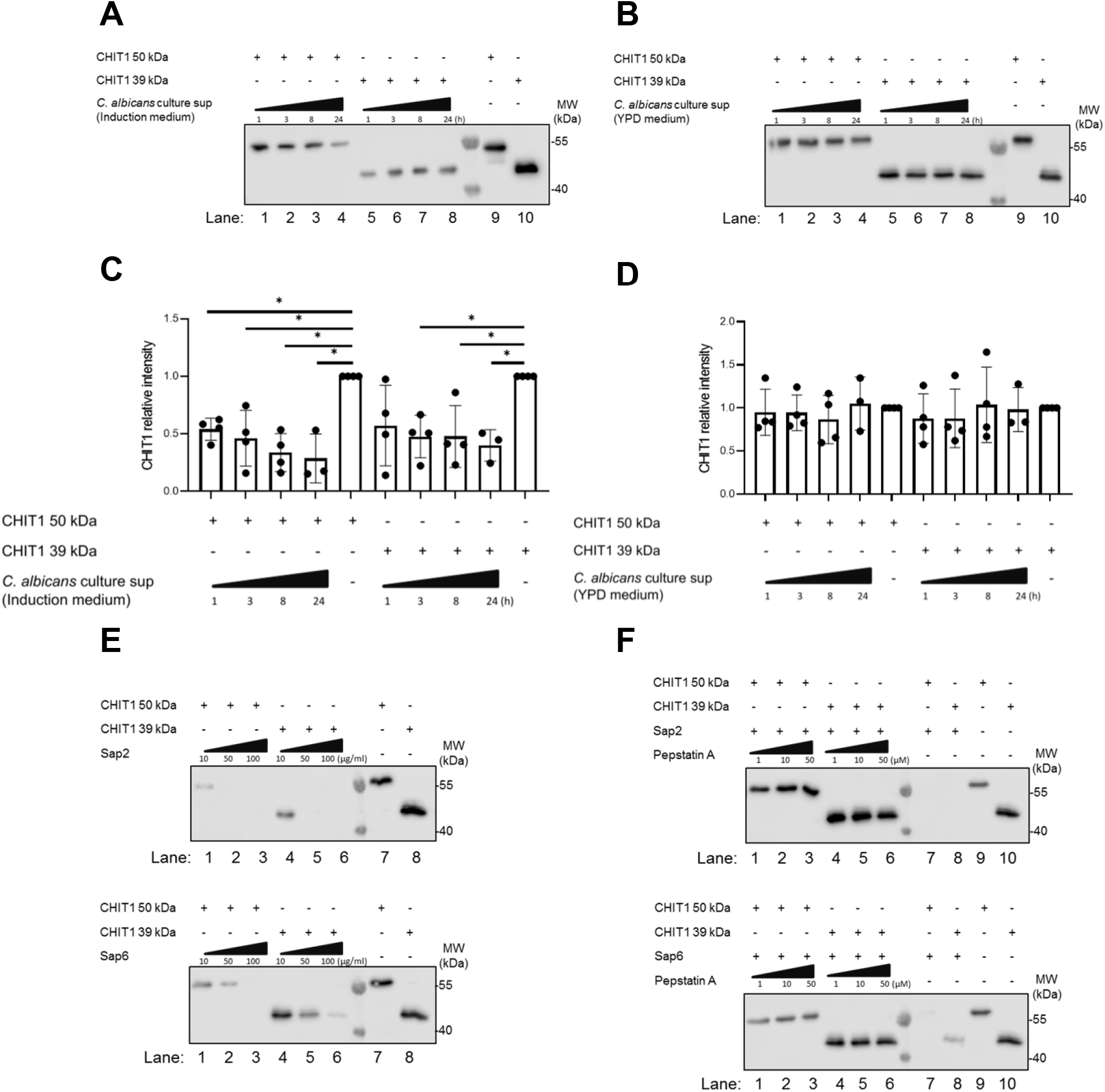
CHIT1 is degraded by secreted *C. albicans* proteases. (A and B) *C. albicans* was cultured for 48 h in protease induction medium (A) or YPD medium (B). *C. albicans* culture supernatants were then collected and incubated with His-tagged variants of CHIT1 50 kDa or 39 kDa for the indicated time points. Recombinant CHIT1 isoforms (His-tagged) were detected by immunoblot with an anti-His antibody. CHIT1 isoforms without any treatment were used as positive controls. (C and D). CHIT1 anti-His relative intensities were quantified and normalized to untreated condition. (E) Recombinant CHIT1 isoforms (His-tagged) were treated with or without the indicated doses of recombinant Sap2 (upper panel) or Sap6 (lower panel) for 8 h and detected as in (A). (F) as in E but with 50 µg/ml Sap2 or Sap6 with or without the indicated doses of pepstatin A aspartic protease inhibitor. In A, B, C (n=4 each) and F (n=2) one representative of ‘n’ biological replicates is shown (mean+SD for technical replicates). C and D (n=4 each) represent combined data (mean+SD) from ‘n’ biological replicates (each dot represents one biological replicate). * p<0.05 according to one-way ANOVA with Dunnett’s correction for multiple testing (C, D).

## Discussion

Here, we uncover that the immune sensing of diffusible chitin oligomers occurs in a much wider context than previously thought. In addition to direct, contact-mediated sensing of chitin MAMP structures by TLR2, our data highlight CHIT1 as a host factor required to generate TLR2-cognate MAMPs, and LBP and CD14 as important components of this sensing pathway (Fig. 7). Furthermore, our data establish TLR1 as a co-receptor in the sensing of the fungal MAMP chitin. The similarities of this system to TLR4 sensing (reviewed in [34]) are unexpected and more striking than previously envisaged: Firstly, both TLR2 and TLR4 detect hydrophobic MAMPs. Secondly, like TLR4, TLR2 appears to operate within the context of the circulating MAMP ‘extractors’ and ‘shuttles’, LBP [34, 35] and CD14 [47, 56, 57], probably owing to the hydrophobic nature of LPS and chitin oligomers. LPS can be extracted from LPS-bearing microbes as a monomer [35], whereas chitin requires degradation by CHIT1 to give rise to immunostimulatory MAMPs. Interestingly, this suggests the chitin-Chit1-LBP-CD14-TLR1/TLR2 cascade resembles the pattern recognition mechanism in *Drosophila* more closely than the LPS-LBP-CD14-MD2-TLR4 pathway in mammals. In *Drosophila*, certain soluble peptidoglycan recognition proteins (PGRPs), the main sensors for bacteria in flies, are hydrolases degrading peptidoglycan. Depending on the receptor involved, peptidoglycan degradation was shown to downregulate or induce host response [58, 59]. Gram-negative binding protein 1 (GNBP1) can also hydrolyze peptidoglycan for recognition by soluble PGRP-SA [12], triggering an extracellular proteolytic cascade ultimately leading to the cleavage of Spätzle, the *Drosophila* Toll receptor ligand [60]. GNBP1 has also been suggested to sense β-1,3-glucans [61] and would thus fulfil the role of a protein binding polymeric MAMPs in flies, whereas our work identifies CHIT1 as a human MAMP generase for a specific PRR. The similarity also extends to the level of protein domains. GNBPs possess carbohydrate binding and glycoside hydrolase domains. Finally, similarities extend to the plant kingdom, in which chitin-degrading enzymes generate oligomeric chitin ligands [5, 62]. Despite small species-specific differences, our work thus defines a unified concept in fungal recognition, involving circulating MAMP generases and linked PRRs, in plant and animal kingdoms.

**Figure 7:**
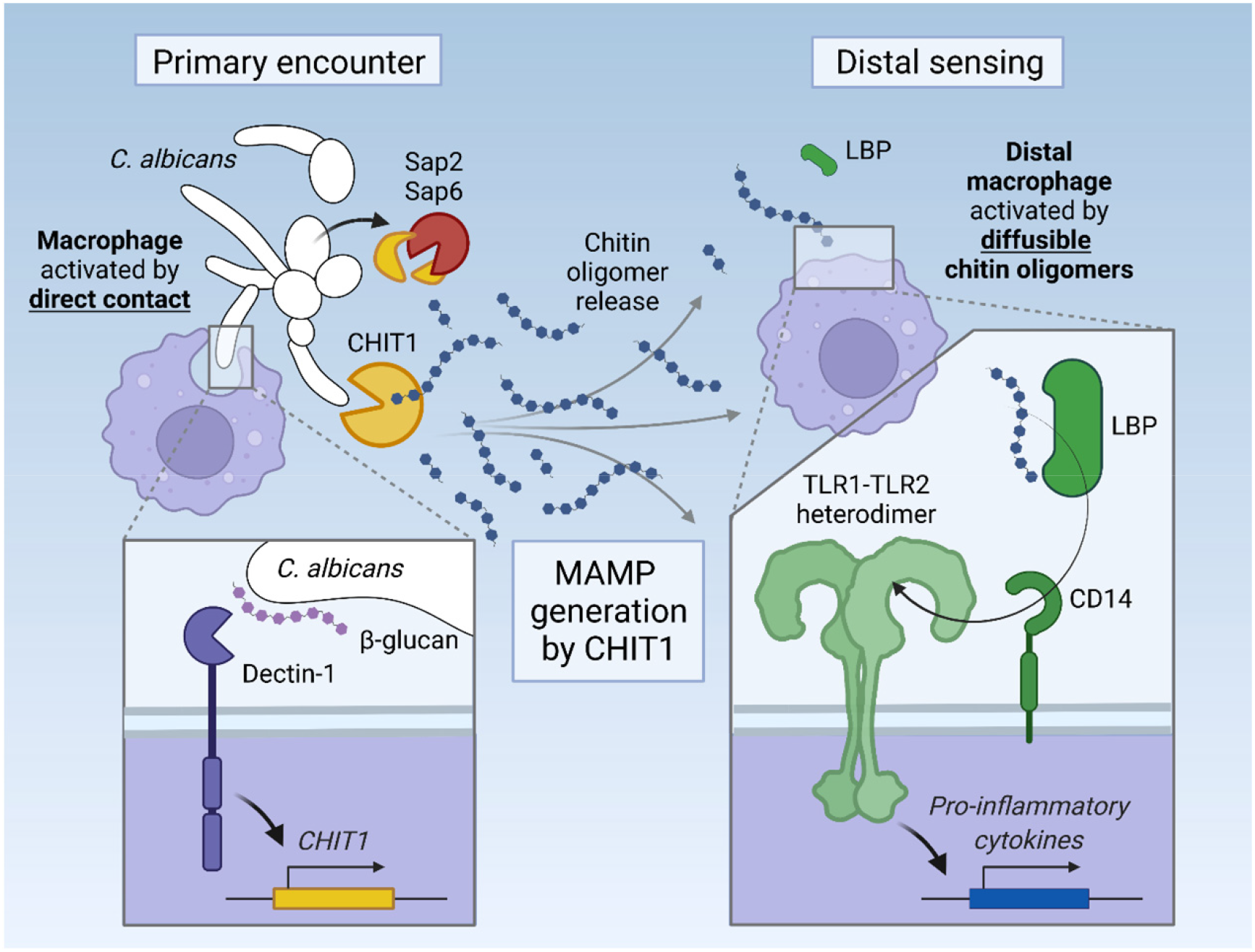
Graphical abstract. Upon its induction via Dectin-1, CHIT1 releases diffusible oligomeric chitin from chitin-bearing organisms which can trigger remote macrophages via an LBP-CD14-TLR1-TLR2 signaling cascade. The activity of CHIT1 may be counteracted by fungal proteases Sap2 and Sap6 which can degrade CHIT1.

The importance of this process for fungal recognition in humans is underscored by evidence that *C. albicans* can target this pathway at the level of MAMP generation through protease activity, reminiscent of counterstrategies used by fungi to subvert immune responses in plants [5]. For example, the plant pathogens *Fusarium oxysporum* and *Ustilago maydis* secrete metalloproteases and/or serine proteases that cleave and thereby inactivate chitinases of tomato and maize, respectively [5]. In mammals, cell wall remodeling exposing chitin may occur as a consequence of changes in carbon availability, pH, exposure to antifungal drug and immune response [63–68]. Targeting circulating CHIT1 via soluble effectors may be an efficient way for pathogenic fungi to avoid subsequent chitin recognition through TLR2. Our findings may explain why no fungal counter-strategies against TLR2 signaling have been described, since targeting the upstream MAMP generase CHIT1 may be favorable. In agreement with our data, patients with *CHIT1* deficiency-associated polymorphisms have higher colonization levels by *C. albicans* [30], and an increased risk of infection by *Madurella mycetomatis*, the causative agent of mycetoma [69]. Our data suggest that impaired TLR2 MAMP generation may contribute to the clinical phenotype of *CHIT1* deficiency in these patients.

Despite the possibility of CHIT1 being targeted by fungi, we suggest that employment of a humoral MAMP generase has advantages for the host over direct recognition of chitin by cell surface receptors. Not only would CHIT1 generate numerous oligomers from a single microbe; the generation of diffusible MAMPs also overcomes the need for innate immune cells to physically engage chitin-bearing microorganisms. For example, phagocytosis and Dectin-1/TLR2-mediated TNF production are critical host responses upon direct engagement of *C. albicans* and other pathogenic fungi [70]. However, the requirement of direct contact would limit responsiveness to locally resident immune or tissue cells and would preclude sensing of MAMPs by distal cells and tissues. Beyond local, contact-based immuno-stimulation, the release of diffusible, oligomeric chitin by CHIT1 enables TLR2 sensing of fungal MAMPs at a distance, providing a means for attraction or recruitment of further immune cells and upregulation of antimicrobial defenses. This might be of particular relevance in epithelia such as the lung, where *CHIT1* is expressed [20] and where a surface area of 180-200 m^2^ [71] requires constant immune surveillance. Here, a sensing system based on diffusible MAMPs is highly advantageous because it requires a limited number of sentinel immune cells. Interestingly, our data also suggest that *CHIT1* expression is induced upon immune sensing of fungi via Dectin-1. This sophisticated mechanism would allow to amplify host innate immune responses by broadening the panel of MAMPs for both local and distal sensing.

Aside from the detection of pathogenic fungi, our data indicate that the link between CHIT1 and the TLR2 sensing system may also be relevant in the context of HDM allergies, where CHIT1-generated chitin oligomers could serve as TLR2-acting adjuvants for HDM antigens. An intriguing link to be explored relates to the Der p 18 allergen from *D. pteronyssinus* : this protein shares a GH18 fold, chitin-binding capability and several conserved GH18 active site residues with CHIT1 [43, 72] and thus might add to oligomer generation. Further analyses of the enzymatic activity and physiological relevance of this and other microbiome-encoded GH18 family members are warranted.

Our results support the notion that CHIT1 enzymatic activity may be an advantageous target for intervention in chitin-TLR2-mediated pathologies such as allergic asthma or Rosacea [73–75]. Interestingly, preclinical studies using CHIT1 inhibitors are under way for the prevention of lung fibrosis ([76, 77]) and might also be efficacious in preventing the generation of inflammatory TLR2 ligands in situ. However, given the abovementioned disparate results obtained in experimental murine in vivo models of *Chit1* deficiency and the differences in *Chit1/CHIT1* expression in mice versus humans [78], an extrapolation from our in vitro data and these mouse models to an application in patients is difficult. However, our data warrant further clinically oriented investigations in this direction.

In conclusion, our data show that CHIT1 is a human MAMP generase for the TLR2 chitin sensing cascade and thereby reveals an astonishingly high level of similarity in how fungal pathogens are sensed in species as different as humans and plants.

## Abbreviations

CHIT1: chitotriosidase
ConA: concanavalin A
ELISA: enzyme-linked immunosorbent assay
HEK: human embryonic kidney
IFN: interferon
iMacs: immortalized macrophages
IL: interleukin
KO: knockout
LPS: lipopolysaccharide
MAMP: microbe-associated molecular pattern
MD-2: myeloid differentiation factor 2
GlcNAc: N-acetylglucosamine
NF-κB: nuclear factor kappa-light-chain-enhancer of activated B-cells
PRRs: pattern recognition receptors
Sap: secreted aspartyl proteases
TLR: Toll-like receptor
TNF: tumor necrosis factor

## Author contributions

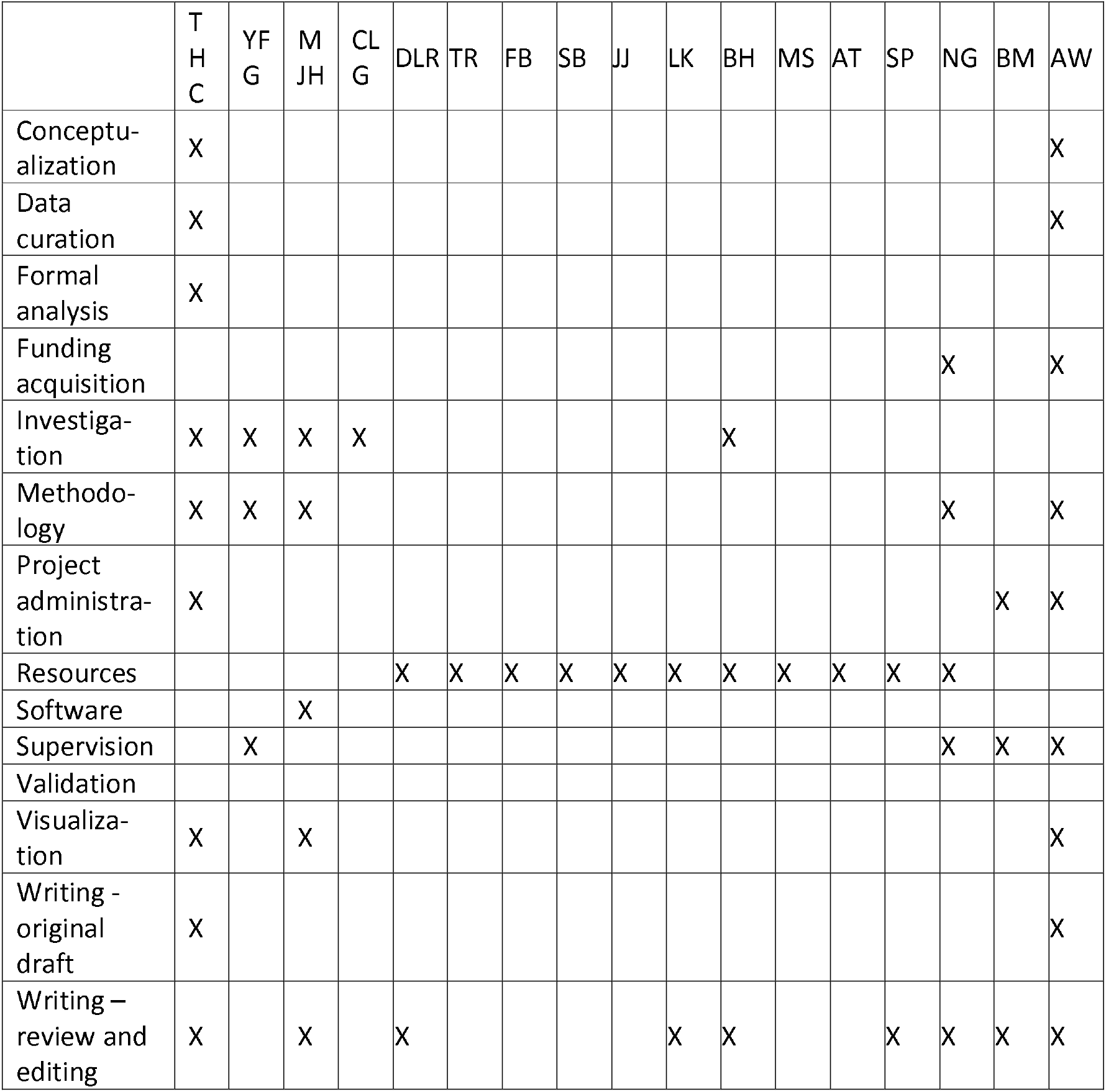

## Acknowledgements

This work was supported by the Wilhelm Schuler Stiftung, the University of Tübingen Medical Faculty, the University of Tübingen Graduate College “Of Plants and Men”, the Deutsche Forschungsgemeinschaft (German Research Foundation, DFG) Collaborative Research Center (CRC) 156 “The skin as a sensor and effector organ orchestrating local and systemic immune responses” (project B05), DFG Priority Program SPP 2225 “EXIT Strategies of Intracellular Pathogens”, DFG Research Grant We-4195/14-1 “Molecular Chitin Sensing by Toll-like Receptors”, and the Federal State Baden-Württemberg Program “Glycobiology”. M.J.H. was supported by a doctoral stipend by the Studienstiftung des Deutschen Volkes. We thank Sabine Dickhöfer and Andrea Dobler for technical assistance and provision of reagents, and Libera Lo Presti for critical reading of the manuscript, editorial support and very helpful comments. TR was supported by the Schweizerischer Nationalfonds zur Förderung der Wissenschaftlichen Forschung (Swiss National Science Foundation grant numbers 320030_149511 and 310030_173123) and by Fondation Carigest/Promex Stiftung für die Forschung (Genève, Switzerland). NG acknowledges Wellcome support of a Senior Investigator (101873/Z/13/Z), Collaborative (200208/A/15/Z, 215599/Z/19/Z), Strategic Awards (097377/Z11/Z) and the MRC Centre for Medical Mycology [MR/N006364/2].

## Materials and methods

### Reagents and the quality control of chitin

All chemicals used in the lab were from Sigma-Aldrich unless otherwise stated. Source and origin of all the ligands, recombinant proteins and inhibitors are listed in Table S1. The preparation of C10-15 chitin oligomers was described before [9]. In brief, C10-15 chitin oligomers were generated from C10-15 chitosan (2000-3000 MW, equivalent to 10-15 subunits, Carbosynth), which was derived from crab shells, chemically hydrolyzed and HPLC-fractionated to >95% purity as confirmed by HLPC and mass spectrometry analysis, see [9] for further details. By using sodium bicarbonate and acetic anhydride acetylation [79], chitosan was acetylated to chitin. The resulting degree of acetylation was assessed upon trifluoracetic acid hydrolysis (for C10-15, 2 h at 100 °C) by ESI and MALDI mass spectrometry, see [9] for further details. Only batches with up to 90% of acetylation were used here. Prior to use for sterile stimulation, acetylated chitin oligomers were suspended in endotoxin free water and tested for endotoxin level by using the limulus amebocyte lysate (LAL) assay (Lonza, CH). Levels below 0.25 EU/ml (<25 pg/ml LPS) in final dilutions were considered acceptable. For levels >0.25 EU/ml, the chitin preparation was incubated for 3 h with 10 µg/ml polymyxin B (Thermo Fisher), washed by centrifugation, and re-assessed.

### Plasmid constructs

The plasmids used in this study are listed in Table S2. Gateway-compatible split-mLumin backbone plasmids for bimolecular complementary assay were described in [80]. Full-length *TLR1* and *TLR2* ORF-containing Gateway Entry plasmids were obtained from the PlasmID Repository at Harvard Medical School. With respect to *TLR6*, a full length ORF flanked with Gateway *attB* sites was synthesized by GENEWIZ and transferred by BP clonase reaction into an *attL* site-containing Gateway Entry plasmid. To transfer the hTLR1, hTLR2 and hTLR6 ORFs into the split-mLumin N-terminal and C-terminal Gateway Destination vectors, respectively, LR reactions using LR Clonase II Enzyme mix kit (Thermo Fisher) were used. The destination vectors were validated by restriction enzyme BsgI (New England Biolabs) (200 – 300 ng plasmid and 5 units enzyme in 1X Tango Buffer, 1 h at 37°C water bath) and Sanger sequencing (GATC Biotech). After sequencing, correct protein expression was verified by Western blotting (see details below). Plasmids encoding the 50 and 39 kDa forms of *CHIT1* as C-terminally His-tagged constructs were described before [42]. To simultaneously introduce the two point mutations D138A and E140L in the catalytic site of the chitinase, the QuickChange II XL site-directed mutagenesis kit (Agilent) was used together with mutagenesis primers (Table S3) designed by following the QuickChange manufacturer’s instruction and verified in silico using Geneious R6 software (6.1.8 version). Presence of the desired mutation and absence of unwanted mutations was confirmed by Sanger sequencing (GATC Biotech). Secretion of the proteins was confirmed by immunoblot and the catalytic inactivity of the mutant CHIT1 proteins was checked by measuring chitinase activity.

### *Candida albicans* maintenance and growth conditions

*C. albicans* strain SC5314 was used in this study, the stock a kind gift from Dr. Anurag Singh (Universitätsklinikum Tübingen), and stored frozen at −80 °C in RPMI medium only (without FCS) containing 20% glycerol. When needed, cells were taken up from the frozen stock and grown at 30 °C in yeast extract-peptone-dextrose (YPD) agar medium (1% [w/v] BactoYeast extract, 2% [w/v] BactoPeptone, 2% [w/v] dextrose, 2% [w/v] agar) on 10 cm dishes. After overnight incubation, cells were stored at 4 °C for up to one month. Before any experiment or treatment, cells were freshly prepared by sub-culturing from 10 cm dish to glass slant tube at 30 °C. Cells were harvested by picking up a smear and re-suspending it in RPMI-1640 medium. To expose the chitin content on the surface of *C. albicans*, yeast cells were incubated with 0.032 µg/ml caspofungin (Sigma-Aldrich) for 6 h at 30 °C in RPMI-1640 medium. After incubation, yeast cells were washed twice with dPBS and tested once for viability. For hyphal induction, 1×10^6^ live yeast cells were re-suspended in YPD medium with 20% FCS and incubated at 37 °C for at least 3.5 h until 90-95% filamentation was observed. The hyphae were collected via a cell scraper and then washed twice with dPBS. For heat-killing, both *C. albicans* yeast and hyphae were prepared by incubation at 65 °C in a water bath for one hour, with killing confirmed by plating in a YPD agar slant tube. For secretion of aspartic proteases (Saps), *C. albicans* was cultured in induction medium (2 mM MgSO_4_, 7.3 mM KH_2_PO_4_, 1% glucose, 0.5% bovine serum albumin, 1% 100× HEPES, pH 4.0) for 48 h at 30 °C as described [55]. The culture supernatants were used to incubate with recombinant CHIT1 and test the degradation effect by immunoblot.

### Recombinant human CHIT1 and *C. albicans* Sap 2 and Sap6

Recombinant 50 kDa and 39 kDa CHIT1 proteins were expressed as C-terminally 8xHis-tagged constructs in HEK293-6E cells and purified from the cell culture supernatant by His-Trap (GE Healthcare) affinity as described in [42]. *C. albicans* Sap2 and Sap6 were expressed in *Pichia pastoris* and purified via ion exchange chromatography and size-exclusion as described in [81].

### House dust mites and chitin flakes preparation

House dust mite *Dermatophagoides pteronyssinus* whole cultures and mite body were from CITEQ Biologics. The powder of whole culture and mite body were weighed and re-suspended in dPBS. Polymyxin B (Thermo Fisher) was then applied at 100 units/ml for working concentration to remove endotoxin. After incubation for at least 1 h at RT, whole culture and mite body were re-suspended in dPBS to 10 mg/ml (stock concentration) and stored at −20 °C. 100 µg/ml of whole culture and mite body were used as working concentration. Chitin flakes from shrimp shell was bought from Sigma-Aldrich (Catalog # C9213-500G). The flakes were firstly sieved by 2 mm or 1 mm pore size steel sieves (Amazon) to sort small pieces of flakes (Fig. S1A). The obtained flakes were picked up by clean forceps into sterile 1.5 ml Eppendorf tubes and treated with polymyxin B (10 µg/ml) for 3 h at RT. The flakes were washed three times with dPBS and finally re-suspended into 1 ml dPBS and stored at 4 °C.

### HEK-Dual^TM^ hTLR2 (NF/IL8) reporter cells

HEK-Dual^TM^ hTLR2 cells were bought from InvivoGen. The cells were derived from human embryonic kidney 293 (HEK 293) and stably transfected with the human *TLR2* gene, an NF-κB/AP-1 inducible secreted embryonic alkaline phosphatase (SEAP) reporter construct and a Lucia luciferase, a secreted luciferase, inserted under the control of the endogenous human*IL8* promoter. Cells were kept under the antibiotic selection of Hygromycin B and Zeocin.

### Immortalized macrophages (iMacs) culture and differentiation

Macrophage-progenitors derived from the bone marrow of C57BL/6 wild-type and *Clec7a*^−/−^ mice were immortalized as described [53] and a kind gift of Philip Taylor, Cardiff University. Progenitor cells were kept and cultured in the RPMI-1640 complete medium supplemented with 1 µM (β-estradiol) and 10% mGM-CSF. Progenitor cells were split 1:10 depending on demand and maintained for less than ten passages. Before differentiation, progenitor cells were washed and centrifuged twice with dPBS to remove β-estradiol. 5 × 10^5^ progenitor cells were then seeded in a 12-well plate with 1.5 ml RPMI-1640 complete medium supplemented with 10% mGM-CSF without β-estradiol. Additional 1 ml of fresh culture medium was added on day 2. On day 5 of differentiation, adherent cells as differentiated macrophages were harvested and plated in 12 well plates (2 × 10^6^ cells/well) in RPMI-1640 complete medium without mGM-CSF. After one day, cells were ready for stimulation.

### N/TERT-1 cell culture

N/TERT-1 cell were a gift from Prof. James Rheinwald [50] and acquired from local collaborator Birgit Schittek, Dermatology Department, University Hospital Tübingen. Cells were passaged and maintained in CnT-07 medium (Epithelial Proliferation Medium, from CELLnTEC, catalog # CnT-07) for less than ten passages. Before stimulation, N/TERT1 cells were seeded at a 96 well plate (2 × 10^4^ cells/well) in CnT-07 medium. After two days of resting, cells were ready for stimulation. After 24 h of stimulation, supernatants were subjected to ELISA according to manufacturer’s instruction (Biolegend).

### Mice and primary bone marrow-derived macrophages

C57BL/6 wild type and *Tlr2* KO mice on a C57BL/6 background (originally a gift from H. Wagner, Ludwigs-Maximilian University, Munich) were maintained in the animal facility of the Department of Immunology, University of Tübingen and used between 8 and 20 weeks of age and sacrificed by using CO_2_. Animal breeding, handling and sacrificing between 8 and 20 weeks of age was performed according to the local institutional guidelines and institutionally approved protocols. WT, *Tlr1* KO, *Tlr2* KO, and *Tlr6* KO mice on a C57BL/6 background were described before [82] and maintained at the animal facility of Lausanne University Hospital, Lausanne, Switzerland. Animal experiments performed in Lausanne were approved by the Service des Affaires Vétérinaires, Direction Générale de l’Agriculture, de la Viticulture et des Affaires Vétérinaires, état de Vaud (Epalinges, Switzerland) under authorizations 876.9 and 3587 and performed according to Swiss and ARRIVE guidelines. Bone marrow cells were isolated from femurs and tibias, which were cut and flushed out by using a 24-gauge syringe, and resuspended in RPMI-1640 medium supplemented with 10% FCS. 1.2 × 10^7^ cells were seeded in 10 cm petri dishes in RPMI-1640 complete medium containing 10% culture supernatant from murine granulocyte-macrophage colony– stimulating factor (mGM-CSF) producing cells [83]. Additional 5 ml of fresh culture medium were added on day 3. On day 7 after differentiation, adherent cells were harvested and plated in 96 well plates (1 × 10^5^ cells/well) or 12 well plates (2 × 10^6^ cells/well) in RPMI-1640 complete medium without mGM-CSF. After one day resting, cells were ready for stimulation.

### Chitotriosidase (CHIT1) digestion

Chitin flakes, *Candida albicans* preparations, house dust mite whole culture and mite body were incubated with human recombinant CHIT1 for digestion. Both recombinant CHIT1 50 kDa and 39 kDa (stock solution 1 µM) were used at a working concentration of 4 nM (1:250 dilution). For expression of WT or mutant CHIT1-encoding plasmids in HEK293T cells, 250 ng of CHIT1 plasmids were transfected into HEK293T cells supplemented with 250 µl DMEM complete medium. After 48 h transfection, the collected cultured medium was used for chitinase digestion. The secretion of both CHIT1 isoforms was confirmed by immunoblot. To digest chitin flakes, *C. albicans* preparations, house dust mite whole culture and mite body were incubated with recombinant or secreted CHIT1-containing media in 1.5 ml Eppendorf tubes on a rotation wheel for 18 h at RT. After CHIT1 digestion, the culture supernatants were used for stimulation or applied to transwells.

### Transwell setting

Transwell inserts for 24-well plate (8 µm pore size, Nunc) were used and 250 µl of media containing chitinase-digested chitin flakes, *C. albicans* or house dust mite bodies filled into the upper reservoir which was placed onto the cell culture plate containing 250 µl media and TLR2-transfected HEK cells or BMDMs at its bottom. After 18 h incubation, almost 2/3 of culture medium from the layer of transwell eventually passed through to the cell culture plate. The transwell was then removed and checked under the microscope. Large particles like chitin flakes and house dust mite were always still trapped in the upper part of the transwell insert. For a schematic impression, see Fig. S1B. For *C. albicans,* minor amounts of yeast cells or hyphae were sometimes found in the lower compartment after incubation. Cells or culture supernatants were processed as described for the respective assays.

### Chitin and chitinase staining on *C. albicans*

Samples containing 2 × 10^6^ *C. albicans* yeast cell or their equivalent of hyphae were transferred to 96-well V-bottom plates. Plates were centrifuged at 5.000 x *g* for 10 min and the supernatants were removed. Cells were resuspended in 100 µl dPBS containing 5 µg/ml of wheat germ agglutinin Alexa Fluor® 647 (WGA, 1:200, Thermo Fisher) and 50 µg/ml concanavalin A Alexa Fluor® 488 (ConA, 1:200, Thermo Fisher), and incubated for 1 h at 4 °C in the dark. For chitinase staining, after incubation with recombinant CHIT1 (His-tagged, [42]), cells were stained with anti-His Alexa Fluor® 594 in 100 µl dPBS overnight at 4 °C (antibody details given in Table S4). Cells were then centrifuged and washed twice with cold dPBS. Cells were resuspended in 5 µl ProLong™ Diamond Antifade Mountant solution (Thermo Fisher). The mounting solution containing stained yeast or hyphae was transferred to glass slides and covered with a coverslip. Slides were stored at RT in the dark until the mounting solution had hardened and then imaged with a Zeiss LSM 800 AiryScan Inverted Confocal Microscope at 630 X magnification in AiryScan mode using a 1.5 X zoom.

### Dual NF-κB luciferase assay in HEK 293 T cells

An inoculum of 5 × 10^4^ HEK 293T cells were seeded in a 24-well plate with 500 µl DMEM complete medium and incubated overnight. The next day, cells were transfected with the following amounts of plasmid DNA per well: 100 ng NF-κB firefly reporter luciferase, 10 ng of *Renilla* luciferase under a constitutive promoter. To measure the TLR2 response, cells were transfected with 100 ng of human TLR2 or with 100 ng backbone as an empty vector control as described [9]. Transfection was performed by using 1 μl of X-treme GENE^TM^ HP DNA Transfection Reagent (Sigma-Aldrich) mixed into total 50 μl Opti-MEM together with the above indicated plasmids. After 15 min of incubation at RT, 50 μl Opti-MEM-X-treme/plasmid mixture were added dropwise onto the cells. After 48 h of incubation, the medium was replaced by fresh DMEM complete medium with or without TLR agonists. Cells were stimulated for 18 h and cell lysates were harvested. The concentrations of all stimuli and ligands are listed in Table S1. To analyze luciferase activity, the culture supernatants were removed and the cells were washed with 350 μl dPBS. Cells were lysed in 60 μl passive lysis buffer (Promega). After shaking for 5 min, cells were frozen in −80 °C for at least 15 min. Thawed lysates were cleared by centrifugation and 10 μl of the cleared lysate was used to measure the luciferase activity on a FluoStar plate reader (BMG Labtech). In the plate reader, the substrates for firefly and *Renilla* luicerase were automatically injected. The analysis settings were used as recommended in the luciferase reporter system by Promega. The results were calculated as the ratio between firefly luminescence and *Renilla* luminescence (Fig. 1A-F, Fig. 4A and H, and Fig. S2).

### HEK-Dual^TM^ hTLR2 (NF/IL8) reporter cells assay

Initially, 5 × 10^4^ HEK-Dual^TM^ hTLR2 cells (Invivogen) cells were seeded in a 96-well plates. After overnight incubation, the culture medium was exchanged with fresh DMEM complete medium with or without the stimuli. After 18 h, cell culture supernatant was analyzed for NF-κB/AP-1-induced SEAP production by using QUANTI-Blue reagents and IL-8-dependent expression of Lucia luciferase using QUANTI-Luc reagents (both Invivogen). For QUANTI-Blue measurement, 20 µl of cell culture supernatant was added to 180 µl QUANTI-Blue solution in a 96-well flat plate. The plate was incubated for 30 min at 37 °C. The SEAP levels were determined by a SpectraMax® plate reader at 650 nm. For QUANTI-Luc measurement, 10 µl of cell culture supernatant was pipetted in a 96-well white plate. Lucia luciferase activity was measured with a FluoStar plate reader (BMG Labtech) which automatically added 50 µl of QUANTI-Luc ™ solution. Data in Figure 1G and 1H were used for this reporter assay.

### Immunoblotting

Protein expression in HEK cells transfected with chitotriosidase- or TLR-encoding plasmids was verified by immunoblotting. In order to detect secreted chitotriosdiases, after 24 h transfection and further 24 h incubation in fresh Opti-MEM without any FCS, both whole cells lysates (WCL) and culture supernatants were harvested. For obtaining WCL, cells were washed once with dPBS and then lyzed with 60 μl RIPA buffer (20 mM Tris-HCl pH 7.4, 150 mM NaCl, 1 mM EDTA, 10% glycerol, 0.1 % SDS, 1 % Triton X-100 and 0.5 % deoxycholate) supplemented with EDTA-free protease inhibitor (Roche) and 0.1 μM PMSF. After maximum speed centrifugation, WCL were mixed with reducing reagent and sample loading buffer (Novex, Thermo Fisher) and then boiled for 5 min at 95 °C to denature the proteins. For obtaining the supernatants, 60 μl of supernatants was treated as described for WCL. Samples (10 μl of WCL and 20 μl of supernatant) were subjected to electrophoresis on 10 % Tris-glycine gels with SDS buffer (25 mM Tris-base, 250 mM glycine and 0.1 % SDS) and transferred to a 0.45 μm nitrocellulose membrane (GE Healthcare). The membrane was blocked with 5 % bovine serum albumin (BSA) (w/v) in Tris-buffered saline solution with 0.1 % (v/v) Tween-20 (TBS-T) for 1 h at RT and then left in 5 ml buffer containing primary antibody (see Table S4) in 5% BSA in TBS-T overnight at 4 °C with rotation. Next day, the membranes were washed three times with TBS-T for at least 5 min and then incubated with 10 ml TBS-T buffer containing secondary antibody in 5% non-fat milk for 1 h at RT with rotation. After incubation with secondary antibodies, membranes were washed three times with TBS-T with at least 5 min per wash. ECL reagent (Peqlab) was used to detect chemiluminescence and the development was performed using Licor camera Odyssey Imaging system in the chemiluminescence channel. Pictures were analyzed and edited in Image Studio^TM^ Lite software. The relative intensities of CHIT1 and –His expression were also quantified by Image Studio^TM^. For the verification of TLR1, TLR2 and TLR6 constructs, TLRs protein expression was verified using by anti-TLR1, -TLR2 and -TLR6 as primary antibodies (Table S4). For whole protein staining, after transfer, nitrocellulose membranes were first rinsed with ddH_2_O then incubated with 5 ml Revert 700 Total Protein Stain Solution (Licor) for 5 min. After washing twice with 5 ml Wash Solution, the membranes were subjected to the Licor Odyssey Imaging system in the 700 nm channel. After imaging, the membranes were destained with 5 ml Revert Destaining Solution for 5 – 10 min until the stain was no longer visible by eyes. The destained membranes were wash twice with ddH_2_O and the 5% BSA blocking was applied immediately.

### Measurement of chitinase activity

Chitinase activity in the culture supernatant of CHIT1 plasmid-transfected HEK293T cells was measured by using a fluorescence assay kit (Sigma-Aldrich Cat. # CS1030). A 10 μl supernatant sample was mixed with 90 μl substrate solution containing 0.5 mg/ml of 4-methylumbelliferyl N,N’-diacetyl-β-D -chitobioside in an assay buffer provided as part of the kit and in a 96-well plate (black/clear bottom plate, Thermo Fisher). The reaction was incubated at 37 °C for 30 min and then stopped by adding 100 μl stop solution. The fluorescence was measured at excitation of 360 nm and emission of 450 nm on a FluoStar plate reader (BMG Labtech).

### ELISA

Appropriate ELISA kits were used to quantify human or murine IL-6 and/or TNF concentrations according to manufacturer’s recommendations (Biolegend). The 450 nm absorbance was measured by a SpectraMax® plate reader.

### Split-mLumin-based bimolecular fluorescence complementary (BiFC) assay

Suspensions of 2 × 10^4^ HEK293T cells were seeded in 8 well 1.5H glass chamber slides (Ibidi). After one day of resting, cells were co-transfected with split-mLumin plasmids in the following two listed combinations: (1). hTLR2-LC151 (200 ng; mLumin C-terminal fragment) + hTLR2-LN151 (200 ng; mLumin N-terminal fragment); (2). hTLR2-LN151 (100 ng; mLumin N-terminal fragment) + hTLR6-LC151 (300 ng; mLumin C-terminal fragment). After 48 h transfection, cells were stimulated with or without C10-15, Pam3 or Pam2 by replacing the culture medium. After 18 h of stimulation, HEK cells were gently washed once with 200 μl dPBS and then fixed with 4% paraformaldehyde in 150 μl dPBS for 10 min at RT. Cells were washed a second time and then stained with Hoechst 33342 (Invitrogen), 1:10,000 dilution in dPBS for 8 min at RT. After the last wash, cells were mounted with Mounting Medium (Ibidi) and imaged by using the Zeiss LSM 800 AiryScan Inverted Confocal Microscope, 400X magnification.

### RT-qPCR analysis

Cultures of 2 × 10^6^ BMDMs were seeded onto 12 well plates after 7 days of differentiation. After one day of resting, cells were stimulated with HDM, *C. albicans* and TLR ligands for the indicated times. Cells were washed with dPBS and lysed in 350 μl RLT buffer (Qiagen) containing 1% (v/v) β-mercaptoethanol, total RNA isolated using the RNeasy Mini kit (Qiagen) following DNA digestion (RNase-free DNase set, Qiagen) and by using a Qiacube instrument. The concentration of total RNA was measured by Nanodrop. mRNA reverse transfection to cDNA was performed by using High Capacity RNA-to-cDNA Kit (Thermo Fisher). Quantitative PCR was performed in 10 μl mix containing 1 to 10 diluted cDNA, TaqMAN Universal MasterMix II and 0.3 μM TaqMan primer and RNA-free water. Each sample was done in triplicates in a real-time cycler (Thermo QuantStudio 7 Flex, Thermo Fisher). The following cycle was: 10 min/ 95 °C; 15 s/ 95 °C and 1 min/60 °C for 40 cycles; cool and save. The primers used are listed in Table S5.

### Fitting of a chitin decamer into a TLR1-2 heterodimer structure

To visualize a chitin decamer bound to a TLR1/TLR2 heterodimer *in silico* (Fig. 4J), docking and an energy minimization simulation were performed as described previously [84]. Briefly, 3D-structures of different GlcNAc oligomers created with the Glycam Carbohydrate Builder (GLYCAM Web, at legacy.glycam.org).were docked into the monomers of the TLR1/TLR2 heterodimer (PDB: 2Z7X, [37]) using AutoDock Vina-Carb 1.1.2 [85]. One docking state each of (GlcNAc)_8_ in TLR1 and (GlcNAc)_10_ in TLR2 were selected and combined via PyMOL (Schrödinger) by aligning and pair fitting to create a TLR1/TLR2 heterodimer with a bound chitin decamer. This new complex is based on the mentioned crystal structure but features a slightly higher distance between the receptor monomers to allow optimal accommodation of the decamer. The outcome was finally optimized using the GROMACS 2019 [86] package with the force fields Amber ff14SB [87] and GLYCAM06 [88] by running an energy minimization simulation with the preparation and setup described previously [84].

### Statistical analysis and software

Experimental data were analyzed in GraphPad Prism 8.3.1 (GraphPad Software, Inc.). Normal distribution was not formally tested but considered to apply judging from the data distribution. Hence two-tailed Student’s *t* tests, one-way or two-way ANOVA tests were used and adjusted for multiple testing as suggested by the analysis software and as indicated. Microscopy data were obtained using ZEN blue (Zeiss) software and analyzed with ImageJ and Fiji. *P* values < 0.05 were generally considered statistically significant and were denoted by an asterisk throughout the figure legends, even if the actual p values were considerably lower. Schematic images were created with Biorender.com.

## Supplemental information

### >Supplemental tables

**Table S1.** List of TLR agonists, recombinant proteins and inhibitors.

| Ligands | Final concentration | Catalog ID | Origin |
| --- | --- | --- | --- |
| C10-15<br>(oligomeric chitin, generated from chitosan) | 20 μM | OC28900 | Carbosynth, see Methods |
| Zymosan | 100 μg/ml | tlrl-zyn | Invivogen |
| Pam3 | 40 nM to 1 μM | tlrl-pms | Invivogen |
| Pam2 | 40 nM to 1 μM | tlrl-pm2s-1 | Invivogen |
| FSL-1 | 40 nM to 1 μM | tlrl-fsl | Invivogen |
| LPS | 100 ng/ml | tlrl-eklps | Invivogen |
| R848 | 5 μg/ml | tlrl-r848-5 | Invivogen |
| Proteins | Concentration | Catalog ID | Origin |
| rCHIT1 50 kDa, rCHIT1 39 kDa | 1 μM | - | This study, [42] |
| Zymolase | 25 nM | tlrl-zyn | Invivogen |
| hLBP | 5 nM to 25 nM | ab151656 | Abcam |
| Sap2 and Sap6 | 10 – 100 μg/ml | - | [55] |
| Inhibitors | Concentration | Catalog ID | Origin |
| Polymyxin B | 10 μg/ml | 21850029 | Thermo Fisher |
| Caspofungin | 0.032 μg/ml | SML0425 | Sigma-Aldrich |
| Pepstatin A | 1 – 50 μM | 40311360025 | Bachem |

**Table S2.** List of plasmids.

| Insert and source if applicable | Tag | Backbone | Resistance |
| --- | --- | --- | --- |
| Empty | n/a | pcDNA3 | Ampicillin |
| hTLR2 | Flag | pcDNA3 | Ampicillin |
| Empty (for BP reaction) | n/a | pDONR207 | Gentamycin |
| hTLR2 (open for gateway)<br>Harvard Plasmid repository | n/a | pDONR221 | Kanamycin |
| hTLR1 (open for gateway)<br>Harvard Plasmid repository | n/a. | pENTR223 | Spectinomycin |
| hCHIT1 50 kDa [42] | His | pTT5V5H8Q | Ampicillin |
| hCHIT1 39 kDa [42] | His | pTT5V5H8Q | Ampicillin |
| Empty<br>Gateway adapted plasmid [80] was<br>a gift from Prof. Qingming Luo | Myc | pDEST_Myc-LC151 | Ampicillin |
| Empty<br>Gateway® adapted plasmid Ref.<br>PMID: 24367102, original plasmid<br>is a gift by Prof. Qingming Luo | HA | pDEST_HA-LN151 | Ampicillin |
| hTLR2-LC151 | Myc | pDEST_Myc-LC151 | Ampicillin |
| hTLR2-LN151 | HA | pDEST_HA-LN151 | Ampicillin |
| hTLR1-LC151 | Myc | pDEST_Myc-LC151 | Ampicillin |
| hTLR1-LN151 | HA | pDEST_HA-LN151 | Ampicillin |
| attB-hTLR6<br>Genewiz synthesis | N.A. | pUC57 | Kanamycin |
| attL-hTLR6 (open for gateway) | N.A. | pDONR207 | Gentamycin |
| hTLR6-LC151 | Myc | pDEST_Myc-LC151 | Ampicillin |
| hTLR6-LN151 | HA | pDEST_HA-LN151 | Ampicillin |
| hCHT1 (50kDa) D138AE140L | His | pTT5V5H8Q | Ampicillin |
| hCHT2 (39kDa) D138AE140L | His | pTT5V5H8Q | Ampicillin |
| NF-κB firefly luciferase reporter<br>(Promega) | N.A. | pNF-κB | Ampicillin |
| *Renilla* luciferase (Promega) | N.A. | pRL-TK | Ampicillin |

**Table S3.**
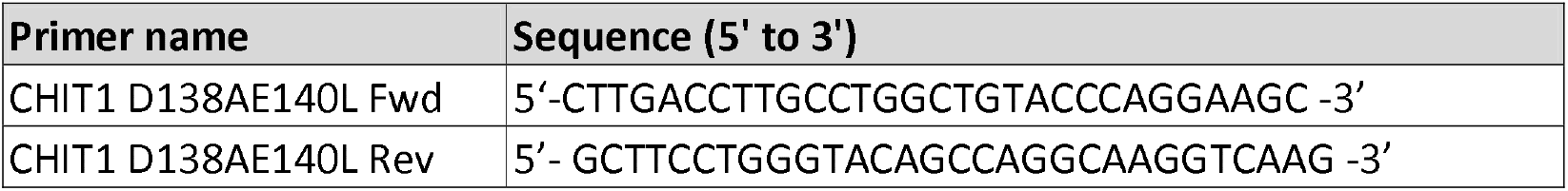
Mutagenesis primers to generate catalytically inactive mutant chitinases.

**Table S4.**
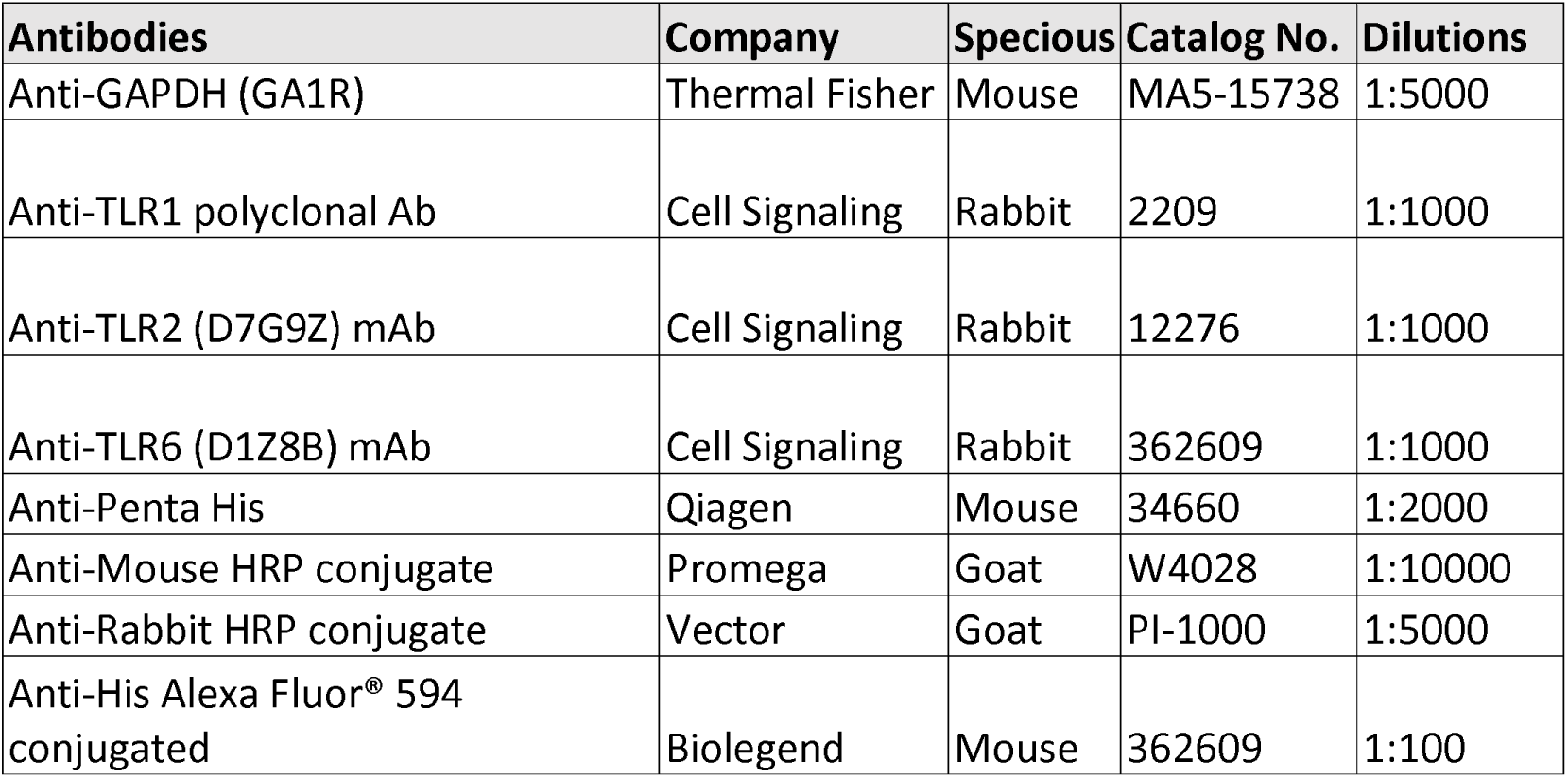
List of antibodies.

| Antibodies | Company | Specious | Catalog No. | Dilutions |
| --- | --- | --- | --- | --- |
| Anti-GAPDH (GA1R) | Thermal Fisher | Mouse | MA5-15738 | 1:5000 |
| Anti-TLR1 polyclonal Ab | Cell Signaling | Rabbit | 2209 | 1:1000 |
| Anti-TLR2 (D7G9Z) mAb | Cell Signaling | Rabbit | 12276 | 1:1000 |
| Anti-TLR6 (D1Z8B) mAb | Cell Signaling | Rabbit | 362609 | 1:1000 |
| Anti-Penta His | Qiagen | Mouse | 34660 | 1:2000 |
| Anti-Mouse HRP conjugate | Promega | Goat | W4028 | 1:10000 |
| Anti-Rabbit HRP conjugate | Vector | Goat | PI-1000 | 1:5000 |
| Anti-His Alexa Fluor® 594 conjugated | Biolegend | Mouse | 362609 | 1:100 |

**Table S5.**
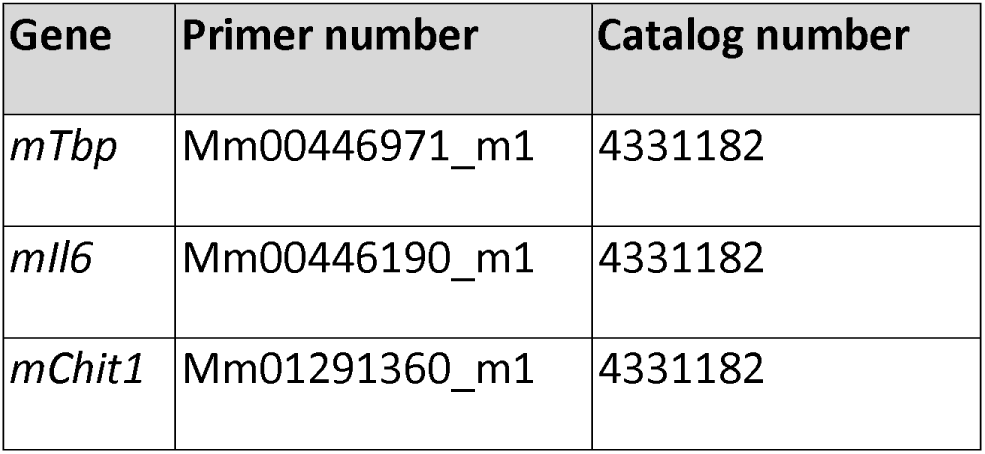
List of qPCR primer.

### Supplemental figures

Figure S1 to S3 uploaded separately. Figure legends included with figures.

