## Supplemental figures for "Transkingdom mechanism of MAMP generation by chitotriosidase (CHIT1) feeds oligomeric chitin from fungal pathogens and allergens into TLR2-mediated innate immune sensing"

### Figure S1

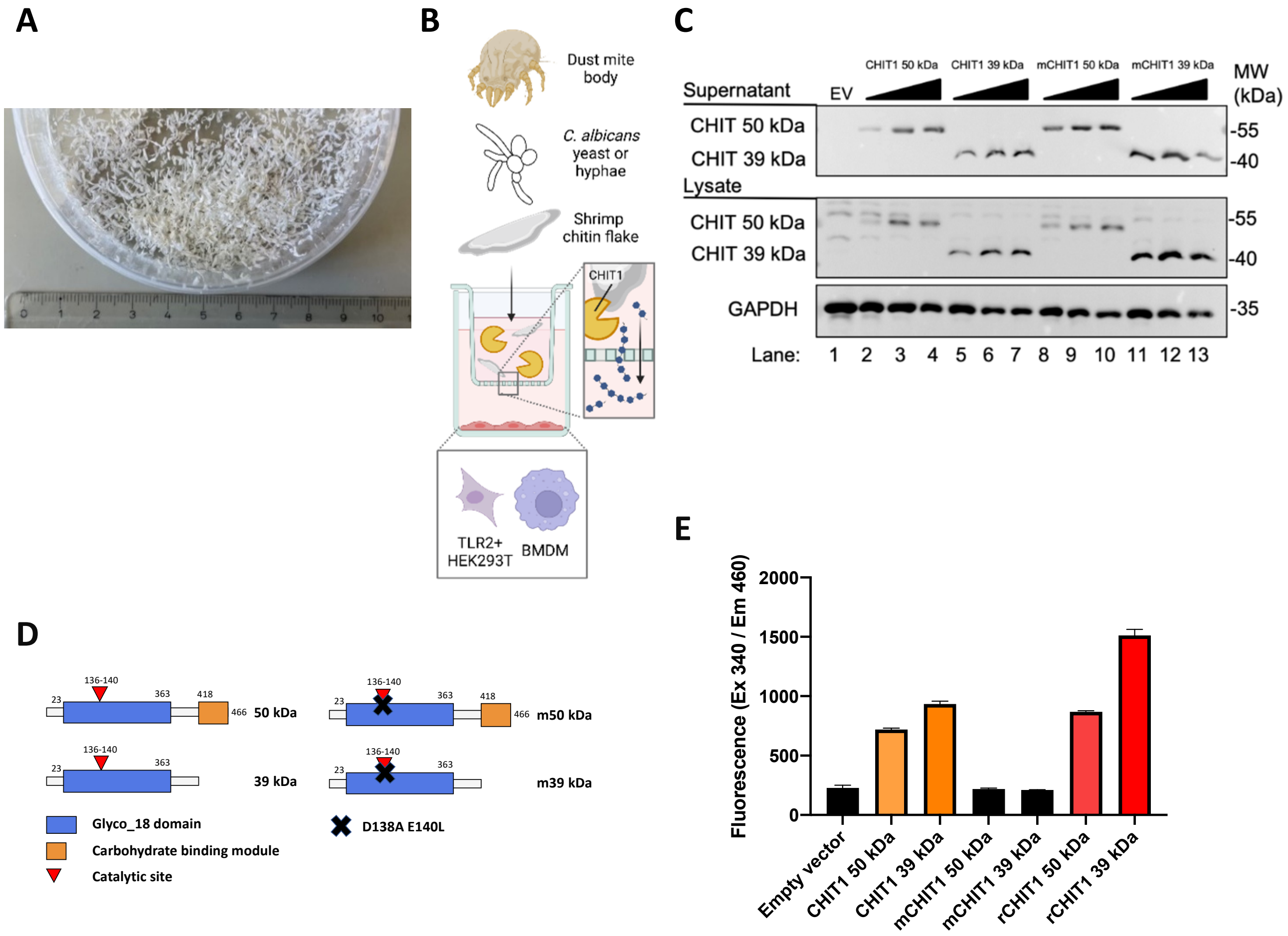


**Figure S1. Catalytic mutant chitotriosidase fails to digest chitin flakes to release diffusible chitin oligomers.**

(A) Size-sorted purified macroscopic chitin flakes from shrimp shells. (B) Transwell setup. Macroscopic stimulants (house dust mite bodies, *C. albicans* yeast or hyphae, shrimp chitin flakes (not to scale) were placed in the upper compartment together with CHIT1. TLR2-expressing HEK293T or BMDM cells in the lower part. (C) Expression of the two isoforms of CHIT1 WT and catalytic mutant protein was assessed by immunoblot with anti-His antibody. (D) Scheme of WT (left) and catalytic mutant (right) CHIT1 isoforms with or without carbohydrate binding module. The black cross indicates the site of introduction of the mutant amino acids within the catalytic site. (E) Measurement of chitinase activity by the hydrolysis of 4-Methylumbelliferyl N,N'-diacetyl-b-D-chitobioside (4-MU-DAC) releasing fluorescent 4-MU. In A, C and E? (n=2 each) one representative of 'n' biological replicates is shown (mean±SD for technical replicates).

### Figure S2

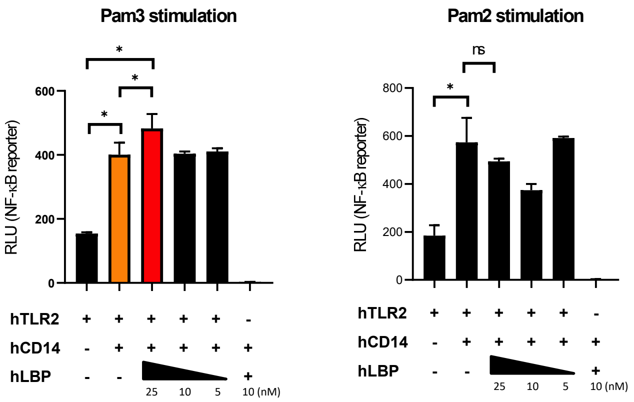


**Figure S2. LBP and CD14 enhance TLR2-NF-κB response to Pam3.**
DLA measurement of NF-κB response in TLR2 or TLR2/CD14 co-transfected HEK 293T cells after stimulation with Pam3 and Pam2. Dose titrated recombinant LBP protein was added together with the stimuli as indicated. Cell lysates were used to measure firefly and Renilla luminescence as described in Figure 1 (C-H). In both panels (n=3 each) one representative of 'n' biological replicates is shown (mean±SD for technical replicates). * $p<0.05$ according to one-way ANOVA with Sidak's correction for multiple testing.

### Figure S3

A

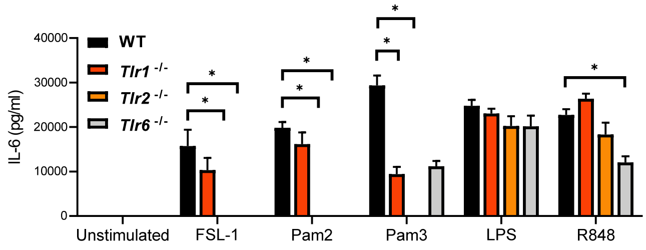


B

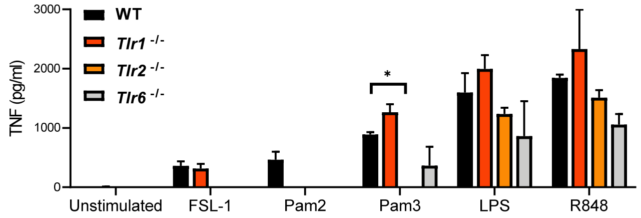


**Figure S3. TLRs receptor specificity to cognate ligands**

(A, B) IL-6 (A) and TNF (B) secretion in murine WT, *Tlr1* KO, *Tlr2* KO and *Tlr6* KO BMDMs was measured via ELISA. FSL-1, Pam2, Pam3 LPS and R848 were used as control ligands to check receptor specificity. In A and B (n=4 each) one representative of 'n' biological replicates is shown (mean±SD for technical replicates). * $p<0.05$ according to two-way ANOVA with Sidak's correction for multiple testing.

#### Figure S4

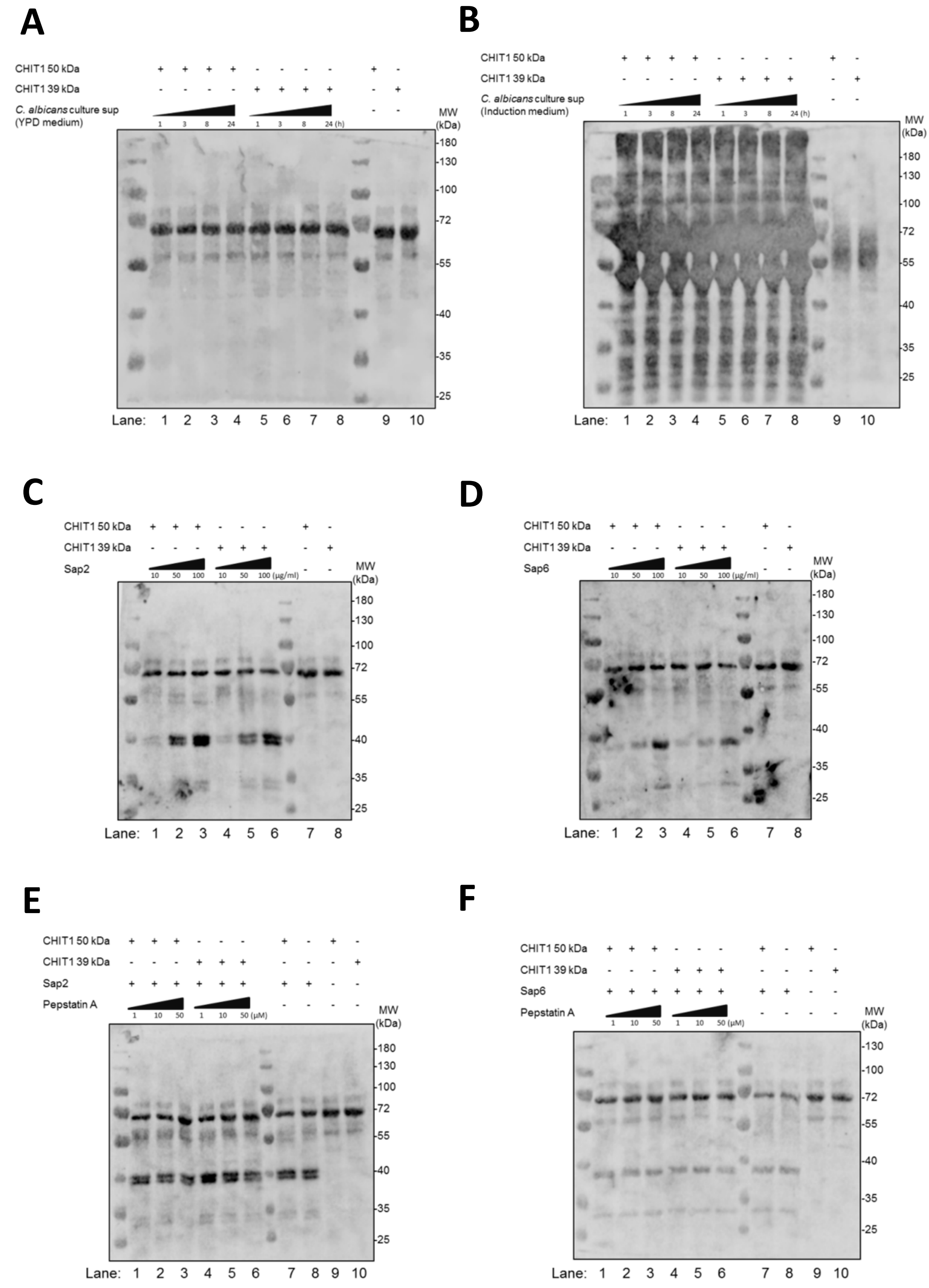


**Figure S4. Whole protein staining of the blots shown in figure 6.**

After the transfer, nitrocellulose membrane was stained with Revert™ 700 Total Protein Stain (Licor) and then subject to imaging in the channel 700 nm by Odyssey® Imaging System. The results of total protein staining: (A) is Fig. 6A, (B) is Fig. 6B, (C) is Fig. 6E upper panel, (D) is Fig 6E lower panel, (E) is Fig 6F upper panel, (F) is Fig 6F lower panel. In A-D (n=3 each), E-F (n=2 each) one representative of 'n' biological replicates is shown.
